# Posterior parietal cortex guides sensorimotor associative learning by linking sensation to distal action

**DOI:** 10.64898/2026.04.19.719425

**Authors:** Qiyu Zhu, Yu Wang, Yuchao Huang, Jieyu Zhang, Xinyi Yang, Angcheng Li, Sixuan Li, Zengcai V. Guo

**Author notes:** These authors contributed equally (Qiyu Zhu, Yu Wang, Yuchao Huang). Correspondence (Z.V.G.).

## Abstract

Sensorimotor associative learning is challenged when sensory cues and motor responses are separated by a delay, limiting the coactivity required for Hebbian plasticity. Here we identify the rostro-lateral posterior parietal cortex (PPC-rl) as a cortical hub that bridges tactile stimuli and delayed licking during learning. Using cortex-wide calcium imaging with single-cell resolution to track ∼16,000 neurons simultaneously across sensory, motor, and association cortices, we find that PPC-rl exhibits sustained delay-epoch activity early in learning that diminishes with expertise. Optogenetic silencing slows learning without impairing expert motor execution. Learning strengthens intra– and inter-areal sensorimotor coupling, and PPC-rl mediates this process by amplifying a low-dimensional communication subspace that aligns sensorimotor co-fluctuations. A biologically plausible network model shows that PPC-rl persistence, subspace alignment, and reward-gated Hebbian plasticity synergize to support delayed association. These findings uncover a PPC-rl based cortical mechanism that maintains temporal continuity to link sensation with distal action during learning.

## Introduction

Sensorimotor associative learning is a fundamental process through which animals learn to link specific sensory cues to appropriate motor responses^1,2^. This type of learning ranges from simple reactions like pressing a button when a light flashes or turning a doorknob in response to a doorbell ring, to more intricate tasks such as playing a musical instrument following a conductor. Between the sensory stimuli and motor action, there is typically a delay, a crucial period for the nervous system to accumulate evidence, retrieve memories, guide decision-making, and prepare future actions^3–6^. Meanwhile, this temporal disconnect also creates the “distal reward problem”: how does an organism associate a delayed action with the cue that caused it, as the neural activity driven by the cue has largely ceased by the time of action that retrieves reward^7,8^?

Classical frameworks of associative learning emphasize activity-dependent synaptic plasticity driven by the concurrent activation of sensory and motor pathways^9–12^. Yet, when sensory stimuli and motor actions are separated by a temporal gap, neurons encoding the cue and those driving the action rarely fire synchronously (Extended Data Fig. 1a-d). Although mechanisms such as neuromodulator-gated eligibility traces have been proposed to preserve transient “synaptic memory” of prior activity^8,13^, these accounts primarily address synaptic timing and do not explain how distributed cortical circuits maintain, route, and transform cue-related information across seconds during learning. Thus, how circuit-level dynamics bridge macroscopic temporal gaps in delayed sensorimotor learning remains unresolved.

To span this temporal divide, the brain could rely on working memory—a process characterized by persistent and transient neural dynamics that maintain cue-specific information through the delay^14–17^. In well-trained animals, working memory representations are widely distributed across sensory, parietal, frontal and motor cortices^5,6^. Coordinated inter-areal communication subspaces align low-dimensional population activity across these regions, creating a delay-spanning channel that preserves the geometry of cue representations and enables downstream regions to selectively transform them into action^18–20^. However, these mechanisms have predominantly been characterized in expert animals, leaving their roles in bridging cues to delayed actions during learning unclear. Moreover, widefield mesoscopic imaging has been used to track cortical average activity changes during sensorimotor learning^21–25^. While these studies have provided valuable insights into global network dynamics, the lacking of single-cell resolution prevents dissection of how distinct functional populations evolve and coordinate across areas during learning. Thus, the central question remains: during learning, how do specific neuronal populations across cortex circumvent the temporal disconnect to forge enduring associations between sensation and distal action?

We hypothesize that an association cortical region outside primary sensory and motor areas sustains delay-epoch activity to bridge sensory cues and distal actions during learning. To test this, we developed a cortex-wide, single-cell-resolution imaging pipeline that simultaneously tracked the activity of ∼16,000 neurons covering sensory, motor, and association cortices throughout the course of sensorimotor learning. Among all regions examined, only PPC-rl exhibited robust delay-period activity that was prominent early in learning and declined as association stabilized. Combining optogenetic silencing with simultaneous cortex-wide imaging to monitor network-level effects, together with computational modeling, we demonstrate that PPC-rl delay activity functionally coordinates somatosensory and motor circuits to facilitate learning. Network simulations suggest delay-activity in sensorimotor cortex as a key driver of associative acquisition and predict that enhancing PPC-rl delay signals accelerates learning—predictions we confirmed experimentally through targeted inhibition and activation. Together, these findings identify the posterior parietal cortex that provides an instructive, delay-specific signal that orchestrates distributed cortical dynamics to bridge sensation and distal action during learning.

## Cortex-wide imaging of neural dynamics during sensorimotor associative learning

To investigate how the brain bridges temporally disconnected sensory and motor events, we developed a delayed tactile detection task in head-fixed mice^26^ (Fig. 1a-b). On the detection trial, a pole vibrated whiskers during the sample epoch (1 s) and mice consumed a drop of liquid milk (16 µl during reward epoch) after a temporal delay (1.5 s). Catch trials had the same temporal structure but omitted whisker stimulation and reward. Before associative training, mice underwent three shaping sessions in which whisker stimuli and milk were delivered at random times, enabling habituation to sensory input and practice of licking without establishing a temporal contingency. Mice learned the task rapidly, reaching stable performance in 4.40 ± 0.24 sessions (mean ± SEM, n = 5 mice, Fig. 1e). Learning was partitioned into three discrete phases (i.e. beginner, intermediate, and expert) based on preparatory licking during the delay epoch (Fig. 1c-d). This classification was concordant with task performance measures (Extended Data Fig. 1e-j) and mirrored changes in orofacial motion energy (Fig. 1e). Beginners licked variably and predominantly after the response cue, whereas experts exhibited anticipatory licking that began during the sample or delay epoch. This behavioral shift highlights the transition from reactive to predictive motor strategies as the association formed.

**Fig. 1.**
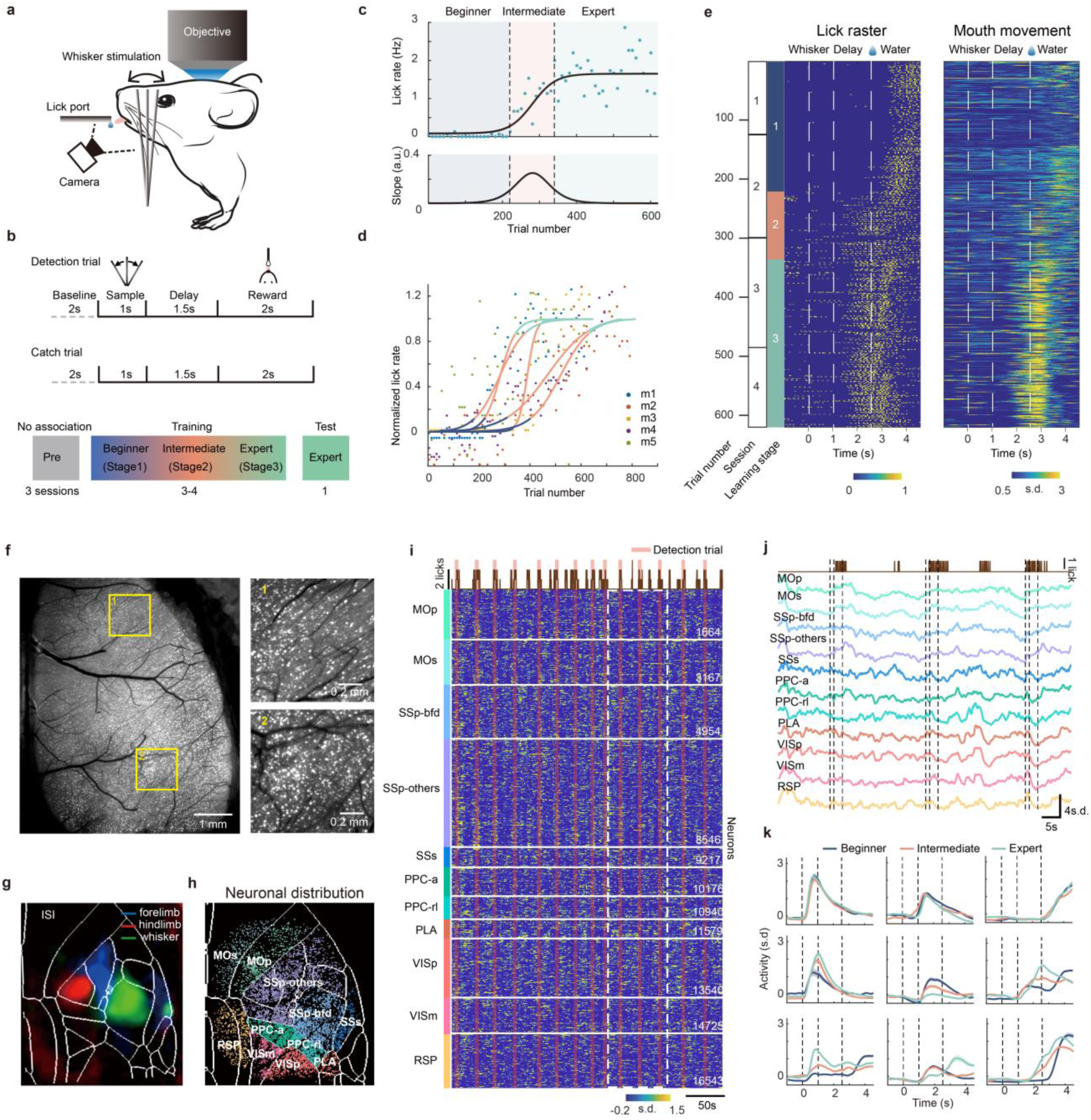
| Cortex-wide single-cell-resolution imaging during sensorimotor associative learning with a temporal delay. **a-e**. Experimental setup and a tactile-based sensory detection task. **a.** Schematic of cortex-wide imaging during learning of the sensory detection task. **b.** Task structure. In detection trials, mice learn to lick after whisker vibrations to consume water reward (top). Bottom: training stages. **c.** Learning stage delineation for an example mouse based on preparatory licking rate in the delay epoch. **d.** Learning stage trajectories for individual mice. **e.** Example behavior across stages from one mouse. Left: lick raster (yellow ticks) for each trial. Right: z-scored mouth-movement traces. **f-k**. Cortex-wide imaging during learning. **f.** Fluorescence showing L2/3 neurons (pixel-wise standard deviation, s.d.). Right, zoom in view of ROIs marked on the left. **g.** Intrinsic signal imaging (ISI) maps for forelimb, hindlimb, and whisker stimulation, overlaid on the Allen Reference Atlas (CCFV3). **h.** Registration of imaged neurons to the atlas, illustrating coverage of MOs, MOp, SSp, SSs, RSP, VIS, PLA and PPC. **i.** Simultaneous calcium imaging of 16,543 neurons across dorsal cortex throughout training. Top, lick rate. Pink shading, duration of detection trials. Traces are z-scored per neuron. Dashed rectangle indicates the window average in **j**. **j.** Mean activity traces for dorsal cortical areas. Vertical lines separate behavioral epochs. **k.** Activity from nine example neurons tracked across learning stages. Mean ± SEM.

To screen for areas underlying sensorimotor learning, we developed a high-throughput imaging pipeline that tracked neuronal activity across the entire dorsal cortex at cellular resolution (Fig. 1f-h and Extended Data Fig. 2a-f and Video 1). We recorded somatic calcium dynamics from layer 2/3 pyramidal neurons using a widefield macroscope (Extended Data Fig. 2a-c, Methods). To circumvent the spatial resolution limits of one-photon imaging in scattering tissue, we optimized the sparsity of labeling to achieve consistent cortex-wide expression, producing spatially isolated fluorescent puncta corresponding to single neurons (Fig. 1f). Two-photon validation confirmed that 86% of widefield-identified cells were resolvable within corresponding two-photon imaging volumes (Extended Data Fig. 2d-f). To enable precise registration of extracted cell coordinates to the Allen Brain Atlas, we additionally performed intrinsic signal imaging (ISI) to map somatotopic boundaries of whisker, forelimb, and hindlimb representations within primary somatosensory cortex (Fig. 1g,h).

In a typical imaging session, we simultaneously imaged 16,543 neurons distributed across somatosensory (whisker SSp-bfd, second somatosensory SSs, and other fields SSp-others), motor (MOp, MOs), posterior parietal (PPC), visual (VISp, VISm), and retrosplenial (RSP) cortices (Fig. 1i,j). Across experiments, we identified 79,314 neurons, with an average number of 15,863 ± 731 (mean ± SEM, n = 5 mice). Neuronal registration across multiple days enabled longitudinal tracking of the same set of neurons across learning (Extended Data Fig. 2g-j). Individual neurons exhibited heterogeneous learning trajectories (Fig. 1k). While subsets of neurons displayed temporally restricted activity modulation (stable, enhanced, or suppressed) within a single epoch (sample, delay, or reward), others dynamically reconfigured their activity across multiple epochs as learning progressed.

## PPC-rl exhibits unique learning-induced neural dynamics

Leveraging cortex-wide single-cell resolution imaging, we tracked learning induced neural dynamics across the dorsal cortex and learning stages. We first examined the spatial distribution of activity by pooling all responsive neurons together (Extended Data Fig. 3f). Averaged activity maps revealed broadly distributed activation even during early learning, masking finer reorganization. We therefore analyzed individual neurons classified by their temporal specificity to task epochs: sample neurons (activity during whisker stimulation), delay neurons (delay-epoch activity), reward neurons (reward epoch), and delay-reward neurons (sustained activity spanning both epochs) (Fig. 2a). These neuron ensembles together accounts about 90% of responsive neurons (Extended Data Fig. 3a-b). To disambiguate task-related activity from motor confounds, we identified lick neurons (activity tightly correlated with spontaneous licking during inter trial interval; Fig. 2a,b). Consistent with this classification, delay, delay-reward and reward neurons were not correlated with spontaneous licking (Fig. 2b and Extended Data Fig. 3c-d).

**Fig. 2.**
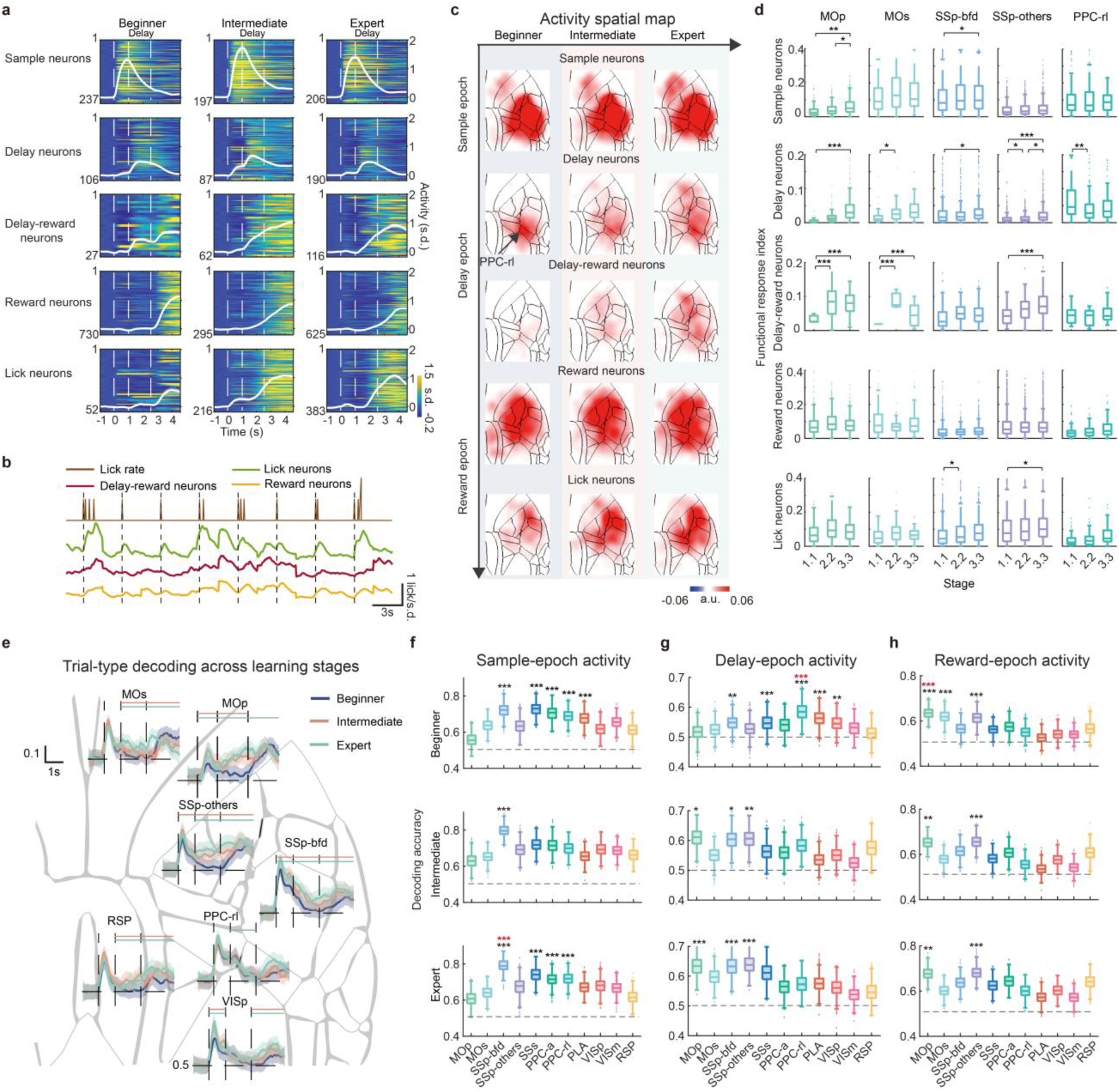
| PPC-rl uniquely sustains delay-epoch activity early in learning. **a.** Activity profiles for each response class of neurons. Heatmaps show single-neuron activity for each response class (sample, delay, delay-reward, lick). Vertical lines separate behavioral epochs. White traces, mean activity for each class (Mean ± SEM). Y-axis numbers, neuron counts. **b.** Only lick-selective neurons exhibit strong correlation with lick rate during ITI. Average activity traces of delay-reward, reward and lick classes from an example animal are shown. **c.** Cortex-wide maps of response-class activity across learning stages. Each row represents one response type. Values are mean activity within the indicated epoch, spatially smoothed with a Gaussian kernel of 10 CCF pixels. Data from 5 mice. **d.** Quantification of response amplitude across areas and learning stages. PPC-rl is the only area with delay-activity declining across learning (for delay neurons). Box and whisker plots show median (line), interquartile range (box), range (whiskers), and outliers (points). Significance is determined by hierarchical bootstrap (\**P* < 0.05, \*\**P* < 0.01, \*\*\**P* < 0.001, Methods). **e.** Trial-type decoding analyses along trial progression based on 300 randomly selected neurons (from all response classes, see Methods). Shading represents SEM estimated using bootstrap. Colored bars indicates the duration with significant difference from beginners (*P* < 0.05, Kruskal-Wallis ANOVA, followed by a post hoc Dunn’s multiple comparison test). **f.** Sample-epoch decoding accuracy by area and learning stage. Top, beginners; middle, intermediates; bottom, experts. Black stars, significant difference from decoding with pooled areas; red stars, significant difference from decoding with pooled data of the stared areas (\*\**P* < 0.01, \*\*\**P* < 0.001, two-sided Wilcoxon rank sum test). **g.** Delay-epoch decoding accuracy as in **f**, highlighting PPC-rl’s elevated contribution early in learning. **h.** Reward-epoch decoding accuracy as in **f**.

We next analyzed the spatiotemporal organization of these neuronal classes and identified four learning-related patterns of cortical reorganization (Fig. 2c-d). First, sample neurons in whisker somatosensory cortex (SSp-bfd) strengthened with learning (indexed as response amplitude weighted by population proportion, Fig. 2d, top row). This observation is consistent with sensory perceptual learning that amplifies task-relevant sensory representations^2,27^. Second, despite no acquisition of novel motor skills, learning increased reciprocal sensorimotor recruitment, with motor cortex activity increased during the sensory epoch and somatosensory cortex activity increased around the licking response (Fig. 2d, top and bottom panels). This mixed sensorimotor information represents a hallmark of sensorimotor integration^26,28–30^. Third, delay and delay-reward neurons in somatosensory and motor cortices surged in activity with learning (130.4% increase from beginner to expert; P < 0.001, Fig. 2c,d, second and third rows). Their activity spanned the gap between stimulus and reward (Fig. 2a), suggesting emergent temporal continuity in sensorimotor circuits. Fourth, in contrast to other regions, delay neurons in PPC-rl peaked during early learning and declined with expertise (intermediate vs beginner mice, 48.87% decrease; P < 0.01, hierarchical bootstrap to account for animals and trials variability; Fig. 2c,d, second row). Notably, among the 11 major cortical areas examined, PPC-rl is the only one exhibiting this significant regression of delay activity (Extended Data Fig. 4a,b).

This bidirectional reorganization, that is the progressive enhancement in sensorimotor cortices and learning-stage dependent regression in PPC-rl, points to region-specific roles. Interestingly, large-scale changes were already evident in the first learning stage when overt behavioral improvements were minimal (substages 1.1–1.3; Extended Data Fig. 3g,h), indicating that coordinated activity reorganization precedes measurable behavioral improvement. We further corroborated the bidirectional reorganization with complementary decoding analyses.

We constructed a pseudo-population for each cortical area by pooling neurons from all response types. We then randomly sampled an equal number of neurons from each pseudo-population and trained a support vector machine (SVM) to classify detection versus catch trials (Fig. 2e-h, Methods). In expert mice, decoding accuracy in whisker somatosensory cortex reached high accuracy (∼80%) with 300 neurons, and we therefore used this fixed population size for all subsequent analyses. Decoder accuracy exhibited a triphasic temporal pattern across trial epochs: peaking during the sample epoch, declining in the delay epoch, and rising again in the reward epoch (Fig. 2e). This temporal pattern was conserved across regions but diverged in magnitude and learning-stage dependency. For decoding using the delay-epoch activity, accuracy in PPC-rl was highest during early learning and declined with expertise (beginner vs. expert, P < 0.01, Kruskal-Wallis ANOVA test, followed by a post hoc Dunn’s multiple comparison test, Fig. 2g), mirroring its activity regression (Fig. 2d). In addition, decoding accuracy was higher in earlier delay compared with the late delay, reflecting pooling of all response types (instead of only the delay group). Decoding accuracy in sensorimotor cortices (SSp, MOp and MOs) showed progressive gains (beginner vs. expert, P < 0.001), aligning with their strengthening delay and delay-reward activity (Fig. 2d,g). For decoding using the sample-epoch activity, the accuracy was highest in the whisker somatosensory cortex (SSp-bfd), with PPC showing intermediate performance at the beginning (Fig. 2f). The accuracy in SSp-bfd further increased with learning, consistent with sensory perceptual learning. Reward-epoch decoding remained stable in sensorimotor areas, reflecting consistent reward and action encoding (Fig. 2h). These results confirm that sensorimotor circuits progressively encode the learned tactile-action association while PPC-rl promotes this temporal linkage in learning, mirroring PPC-rl’s unique learning dynamics.

## PPC-rl plays a causal role in sensorimotor associative learning

Cortex-wide imaging revealed a distinctive pattern of PPC-rl delay activity that was prominent during early learning and declined as mice became proficient. We then asked whether PPC-rl played a causal role for learning. In total, we used 79 mice with optogenetic perturbation or control conditions across learning (Methods). To test at which stage PPC-rl is required, we first bilaterally inhibited PPC-rl on randomly interleaved probe trials using temporally precise optogenetic perturbation (Fig. 3a, Methods). Delay-epoch inhibition did not reduce baseline preparatory licking in beginner mice and had minimal effects in experts (Fig. 3b). However, the inhibition significantly reduced preparatory licking in intermediate mice (34±8.1% vs 3.4±2.7% reduction in experts, mean ± SEM, Fig. 3b), suggesting that PPC-rl might be required during learning but becomes dispensable for well-learned execution.

**Fig. 3.**
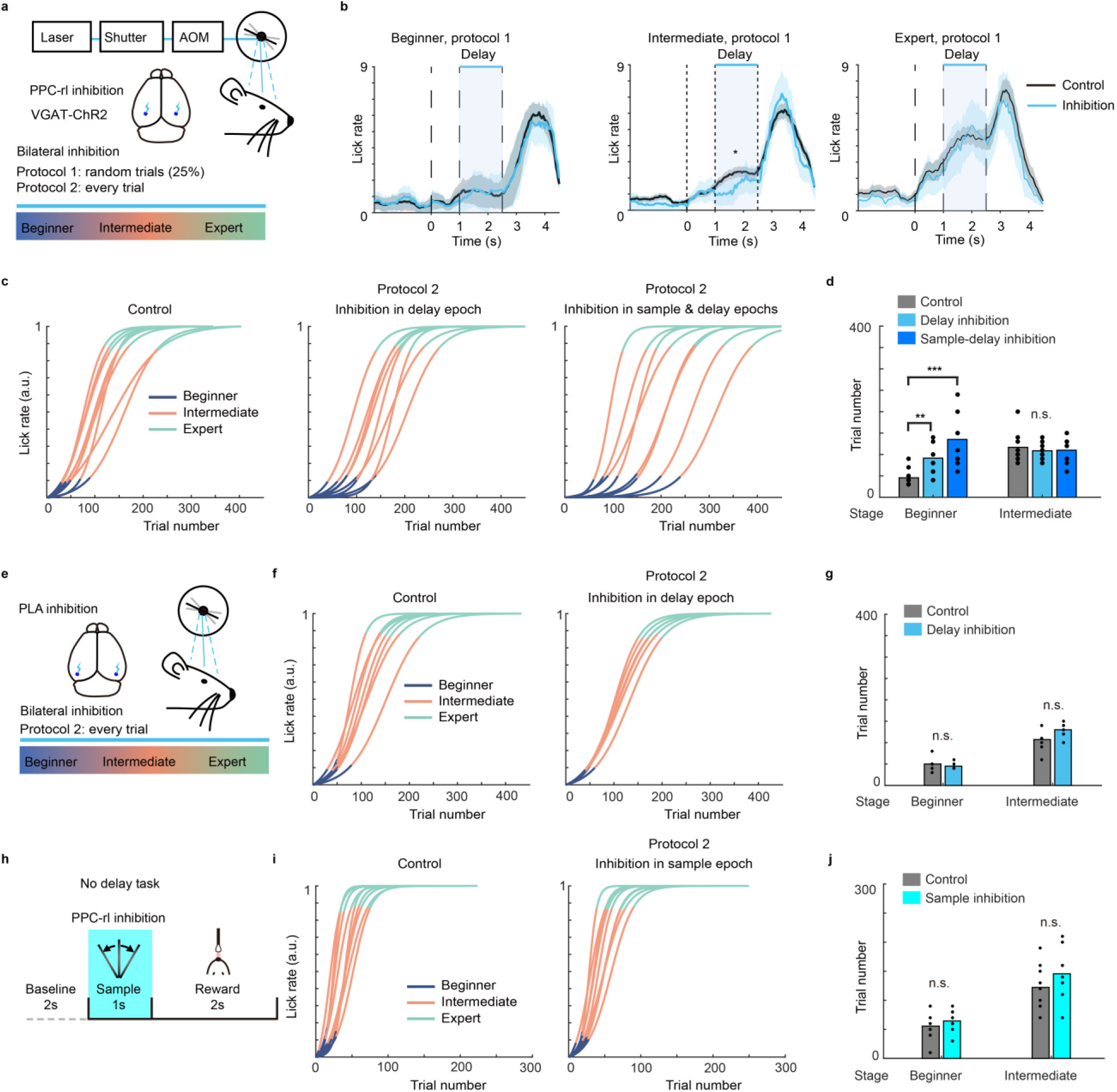
| PPC-rl is required for learning but dispensable for expert sensorimotor execution. **a.** Schematic of optogenetic inhibition. Bilateral PPC-rl was suppressed via scanned laser illumination with a galvanometric mirror system (see Methods). **b.** Delay-epoch inhibition reduced preparatory licking in intermediate mice (left) but not in exerts (right). Inhibition was deployed on 25% of randomly interleaved trials (n = 5). Shading represents SEM. \**P* < 0.05, \*\**P* < 0.01, \*\*\**P* < 0.001, two-sided Wilcoxon rank-sum test for all behavioral comparisons, related to **b, d, g, j**. **c.** PPC-rl inhibition slows acquisition of the tactile-lick association. Left, control mice without inhibition (n = 9 mice; note lines overlapping; mice learned faster compared with imaging mice in a different genetic background, see Methods). Middle, inhibition during the delay epoch (n = 8 mice). Right, inhibition during the sample and delay epochs (n = 8 mice). For each cohort of inhibition mice, inhibition was applied on every trial (**c-d**). Control mice and inhibition mice have the same genetic background. **d.** Quantification of the number of trials in each learning stage, demonstrating prolonged early stage with PPC-rl inhibition. Same data as in **c**. **e.** Schematic of optogenetic inhibition of PLA. **f.** Learning stages during control and PLA inhibition. Same inhibition protocol as PPC-rl. Left, control mice without inhibition (n = 7 mice; mice learned faster compared with imaging mice with a different genetic background, see Methods for discussion). Right, inhibition during the delay epoch (n = 6 mice; note lines overlapping). **g.** Quantification of the number of trials in each learning stage. PLA inhibition does not impair the tactile-lick association. n.s., not significant. Same data as in **f**. **h.** Schematic of the no-delay task and PPC-rl inhibition. **i.** Learning stages during control and PPC-rl inhibition. Left, control mice without inhibition (n = 9 mice; note lines overlapping; mice learned faster compared with imaging mice with a different genetic background, see Methods for discussion). Right, inhibition during the sample epoch (n = 9 mice). **j.** Quantification of the number of trials in each learning stage. PPC-rl inhibition in the no-delay task does not affect tactile-lick association. Same data as in **i**.

To directly test the learning role of PPC-rl, we performed PPC-rl inhibition on every trial throughout training. Mice receiving delay-epoch inhibition completed a comparable number of trials per session as control mice (106.71 ± 3.87 vs 106.18 ± 4.67, respectively; mean ± SEM), and reached similar levels of preparatory licking after learning (5.37 ± 0.31 vs 4.43 ± 0.30, respectively; p > 0.05, two sided Wilcoxon rank-sum test). However, delay-epoch inhibition significantly slowed learning, prolonging time spent in the beginner stage and mice required more trials to reach expert performance (Fig. 3c,d). We then examined the specificity of PPC-rl effect on learning. First, the contribution of PPC-rl is delay-task dependent. When the delay epoch was removed from the sensorimotor association task, PPC-rl inhibition produced minimal behavioral impairment (Fig. 3h-j). Second, the causal role is region specific. Inhibition of the posterior lateral association cortex (PLA)—which also showed delay activity decoding in beginner mice (Fig. 2g)—did not alter learning rate (Fig. 3e-g).

Because PPC-rl delay activity depends on whisker-evoked SSp-bfd input (Extended Data Fig. 4c-g), we additionally inhibited PPC-rl during both the sample and delay epochs. This combined perturbation further impaired learning compared with delay-only inhibition (Fig. 3c,d), consistent with a model in which sensory drive reaches PPC-rl during the sample epoch, and PPC-rl subsequently sustains delay activity to support association formation. In summary, these experiments demonstrate that PPC-rl plays an area-selective, stage-dependent, and delay-task specific role in linking sensation to distal action during learning.

## Learning-driven reorganization of cortex-wide functional connectivity

Linking a tactile cue to delayed licking likely requires learning-dependent reconfiguration of interactions among sensory, association, and motor cortices. To test whether PPC-rl participates in this reconfiguration, we quantified functional connectivity (FC) from noise correlations during inter-trial intervals (ITIs), thereby isolating intrinsic co-fluctuations from task-locked signals. Two features improved robustness of this estimation: 1) long ITIs providing a large number of time points for correlation estimation, 2) simultaneous cortex-wide recordings of ∼16,000 neurons per session, yielding hundreds to thousands of neurons per area. To further minimize behavioral confounds, we excluded movement periods from ITIs, as movement broadly elevates brain-wide activity (Methods)^31–33^.

This dataset enabled internal comparisons at multiple levels. Simultaneous multi-area recording allows regional comparisons (e.g. sensorimotor vs. non-sensorimotor areas), and single-cell resolution enables comparisons across functionally defined populations (e.g. sample vs delay neurons). We first assessed FC among matched response classes across cortical areas (sample, delay, delay-reward, lick; Fig. 4a). Intra-areal FC within a response class remained stably high (∼0.1) across learning (Fig. 4a, diagonal). In contrast, inter-areal FC increased selectively with learning: cross-area coupling strengthened most for delay-reward and lick-related neurons, modestly for delay and reward neurons within sensorimotor regions, and remained weak for sample neurons (Fig. 4a, off-diagonal; Fig. 4d).

**Fig. 4.**
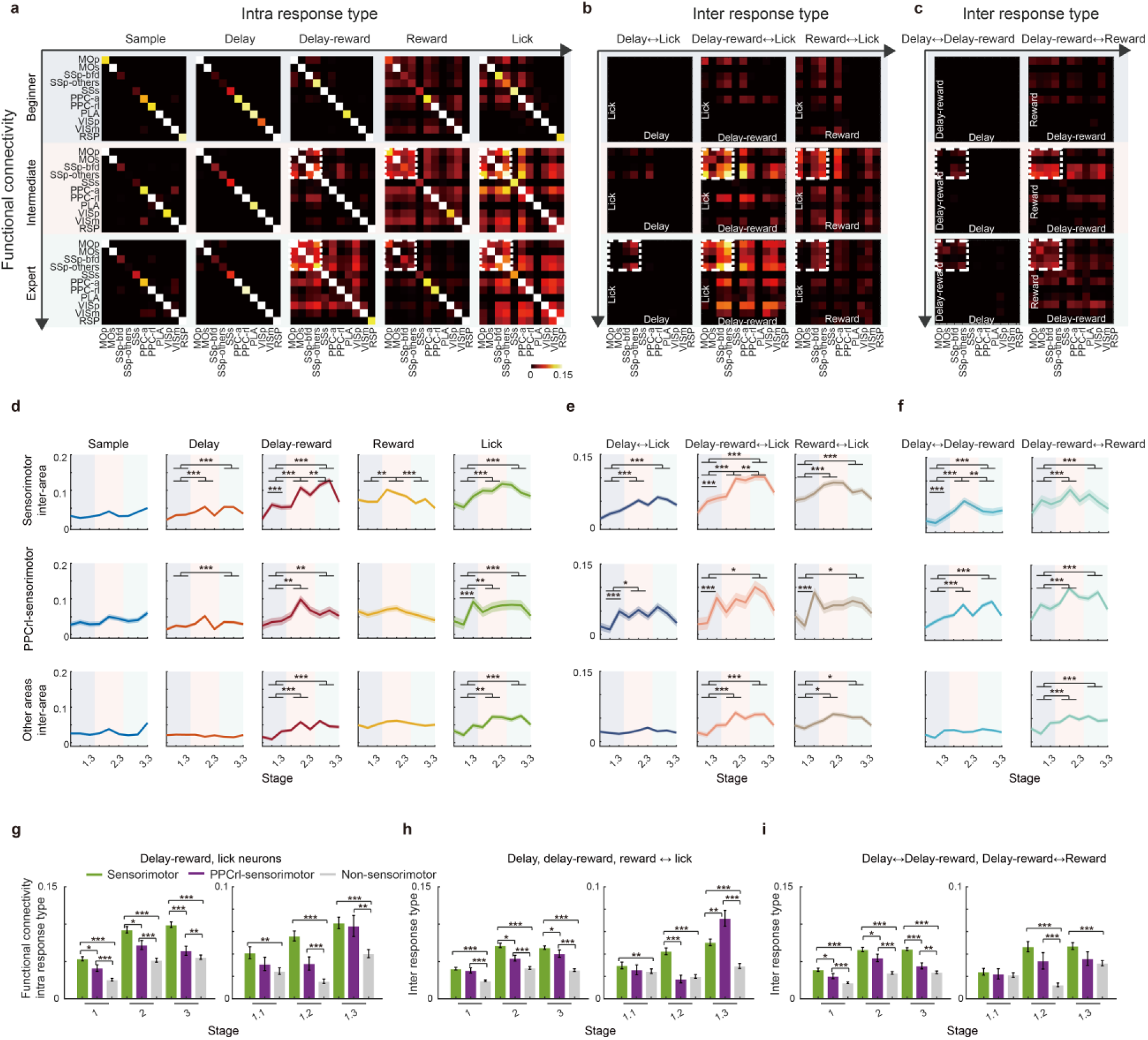
| Learning strengthens selective inter-areal functional coupling. **a.** Functional connectivity (FC) for matched response types (intra-response types: sample, delay, delay-reward, reward, lick). Heatmaps reflect noise correlations (835,441,216 neuron pairs from 5 mice) between indicated cortical areas. Rows, learning stage (top, beginner; middle, intermediate; bottom, expert). Columns, response types. Axes indicate cortical areas. **b.** FC between response types (inter-response types: left, delay↔lick; middle, delay-reward↔lick; right, reward↔lick). Axes denote neuronal response types and cortical areas. **c.** Inter-type FC along the nominal temporal chain (left, delay↔delay-reward; right, delay-reward↔ reward). Same format as in **b**. **d.** Intra-type FC across learning for different areas. Top: inter-areal FC within sensorimotor cortices. Middle: FC between PPC-rl and sensorimotor areas. Bottom: other cortical regions. Columns correspond to response-types. Statistics: \**P* < 0.05, \*\**P* < 0.01, \*\*\**P* < 0.001, Kruskal-Wallis ANOVA test, followed by a post hoc Dunn’s multiple comparison test. Shading, SEM. **e.** Inter-type FC across learning for links involving the lick population (with delay, delay-reward, reward populations). Same format as in **d**. **f.** Inter-type FC across learning for links involving temporal chain pairs (left, delay↔delay-reward; right, delay-reward↔ reward). Same format as in **d**. **g.** Quantification of intra-type FC showing strength difference between different cortical circuits and stages. Left, FC for learning stages 1, 2, and 3 (beginner, intermediate, expert). Right, sub-stages within stage 1 (1.1-1.3). FC from delay-reward and lick neurons are averaged (\**P* < 0.05, \*\**P* < 0.01, \*\*\**P* < 0.001, Kruskal-Wallis ANOVA test, followed by a post hoc Dunn’s multiple comparison test). Error bar, SEM. **h.** Quantification of inter-type FC for lick-targeted links (averages from **b** and **e)**. Same format as **g**. **i.** Quantification of inter-type FC for temporal chain pairs (averages from **c** and **f)**. Same format as **g**.

We then assessed inter-type FC between neuronal groups with distinct temporal selectivity. We first examined putative connections by pooling each class of neurons across the dorsal cortex (Extended Data Fig. 5). Because the learned response is tactile-contingent licking, we focused on putative links from delay, delay-reward, and reward populations to lick neurons. Given the learning-dependent increase of delay and delay-reward activity, we also examined the nominal temporal chains from delay to delay-reward and from delay-reward to reward groups. Links involving lick neurons and the temporal chain pairs were significantly stronger than the global average, and further increased with learning (Extended Data Fig. 5a-c). Granger causality computed from within trial activity revealed significant directed influences among these pairs (Extended Data Fig. 5d-i).

We next examined FC between pairs of cortices and identified areal specificity (Fig. 4b,c,e,f). Within sensorimotor cortices, inter-type FC strengthened markedly with learning for connections involving lick neurons and for the temporal chain pairs. Cross-region FC between PPC-rl and sensorimotor areas increased for delay↔lick, delay-reward↔lick, reward↔lick, and delay↔delay-reward pairs. Outside sensorimotor circuits, only delay-reward↔lick, reward↔lick, and delay-reward↔reward connections strengthened. Across learning, FC within sensorimotor cortices and between PPC-rl and sensorimotor areas remained significantly higher than FC outside these networks (Fig. 4g-i).

These FC changes tracked behavioral improvement. Connections ramped up in parallel with performance gains during stages 2-3 (Fig. 4d-f). In stage 1, inter-areal FC within sensorimotor cortex was stable or only modestly increased (delay-reward↔lick and delay↔delay-reward, increased by 37.6 ± 2.91% from stage 1.1 and 1.2 to 1.3, mean ± SEM). Intriguingly, the PPC-rl-sensorimotor FC (delay↔lick, delay-reward↔lick, reward↔lick) exhibited an early surge at the end of stage 1, preceding overt behavioral improvement (Fig. 4e,f middle row, Fig. 4h, right, 229 ± 3.93% increase mean ± SEM). This temporal lead positions PPC-rl as an early coordinator that primes sensorimotor communication to facilitate association formation. Together, these results indicate that learning strengthens selective, directed functional coupling within sensorimotor circuits and along PPC-rl-sensorimotor pathways.

## PPC-rl promotes co-fluctuation of somatosensory and motor cortical activity during learning

Given PPC-rl’s distinctive delay dynamics and causal contribution, we asked how PPC-rl shaped sensorimotor coupling across learning. We optogenetically inhibited PPC-rl specifically during the delay epoch at defined training stages while simultaneously imaging somatosensory and motor cortices to monitor the inhibition effects (Fig. 5a, Methods). Inhibition on randomly interleaved trials was achieved by illuminating virally expressed eNpHR3.0 in PPC-rl, with the PPC-rl boundary delineated via registration to the Allen Brain Atlas. The registration was anchored by ISI maps of the whisker, forelimb, and hindlimb somatosensory areas (Fig. 5b). We verified inhibition efficacy with three measures. First, we observed strong and spatially localized suppression at the injection site (Fig. 5c and Extended Data Fig. 6b). Second, there was a ring of increased activity surrounding the injection core (Fig. 5c), consistent with disinhibition via cortical lateral interactions^34^. Third, PPC-rl inhibition modestly but significantly reduced preparatory licking during intermediate learning (Fig. 5d,e and Extended Data Fig. 6c,d), aligning with the stage-specific behavioral effect seen with transgenic mice (Fig. 3b).

**Fig. 5.**
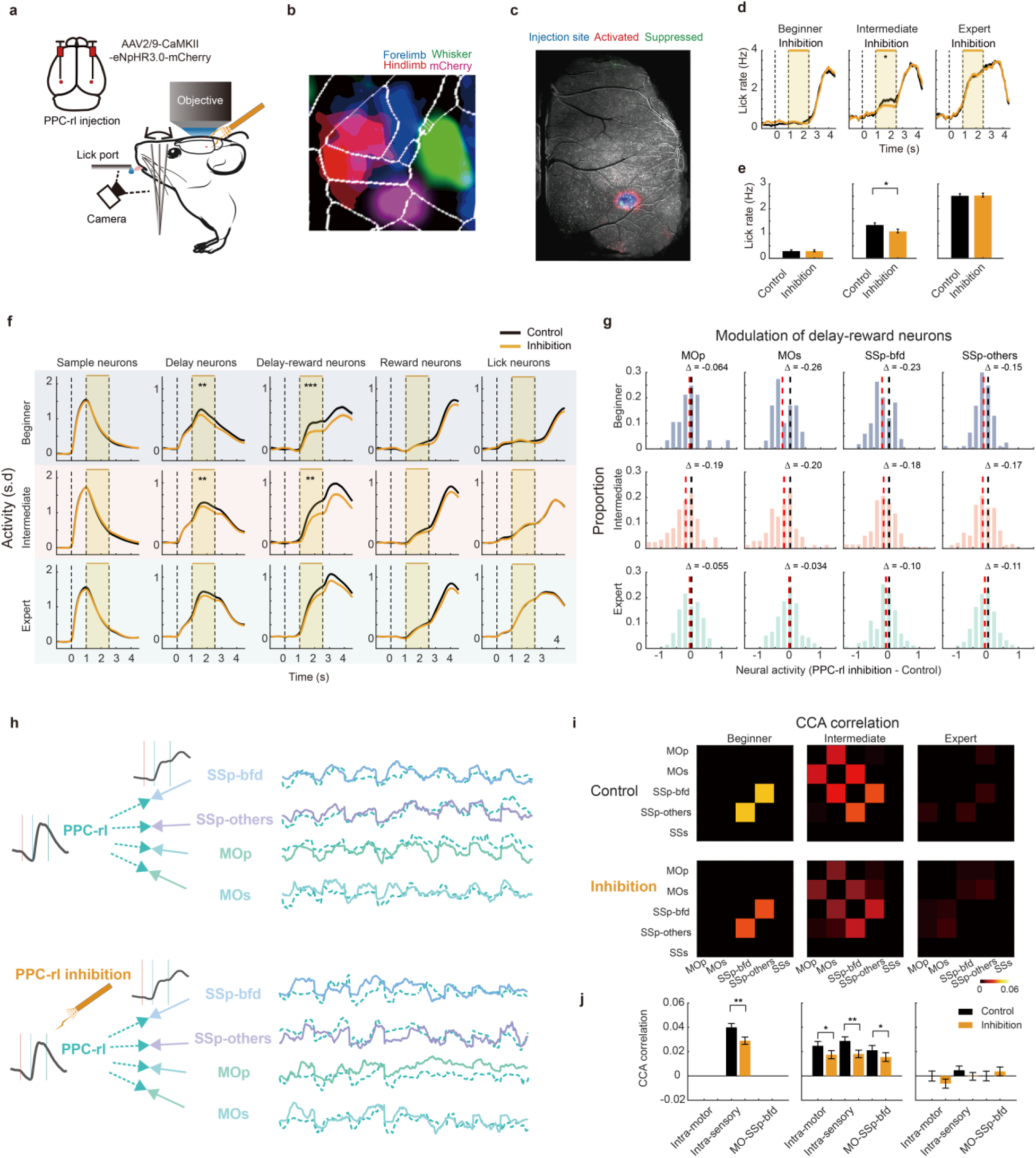
| PPC-rl enhances delay-epoch activity in sensorimotor cortices and drives their co-fluctuations during learning. **a.** Experimental schematic for bilateral PPC-rl inhibition during cortex-wide imaging. AV2/9-CaMKII-eNpHR3.0-mCherry virus is injected in PPC-rl. In the imaged hemisphere, 589 nm laser light is delivered through a collimating lens. In the contralateral hemisphere, laser light targets PPC-rl through an optical fiber. Inhibition is restricted to the delay epoch. **b.** Viral targeting of PPC-rl via ISI. PPC-rl boundary is registered to the Allen atlas, anchored by ISI maps for forelimb, hindlimb, and whisker somatosensory areas. Viral injection is targeted to PPC-rl after boundary delineation. **c.** Imaging field of L2/3 neurons showing pixelwise fluorescence s.d. overlaid with the injection core. Inhibition produced local suppression with a ring of excitation that is outside of PPC-rl. **d.** Delay-epoch PPC-rl inhibition reduced preparatory licking in intermediate mice. Orange, inhibition; black, control. Shading, SEM. Lick rate in the delay epoch was averaged for comparison (\**P* < 0.05, two-sided Wilcoxon rank sum test, n = 3). Related to **d-g, i, j**. **e.** Quantification of inhibition effects on preparatory lick rate across learning stages. Same data as in **d**. **f.** PPC-rl inhibition suppressed delay– and delay-reward activity in beginner and intermediate mice, but not in experts (\*\**P* < 0.01, \*\*\**P* < 0.001, two-sided Wilcoxon rank sum test). Shading, SEM. **g.** Modulation for delay-reward neurons in sensorimotor cortex. Red dashed line, median under PPC-rl inhibition. Black line marks zero. **h.** Canonical correlation analysis (CCA) identifies low-dimensional communication subspaces between PPC-rl and individual sensorimotor areas. **i.** Learning-stage dependent co-fluctuations in the first CCA mode between sensorimotor pairs (e.g., SSp-bfd ↔ SSp-other, MOp ↔ MOs, SSp-bfd ↔ MOs). The co-fluctuations are strongest in beginner and intermediate mice while diminished in experts. **j.** Quantification of CCA-derived coupling across area pairs and learning stages, showing significant inhibition induced reductions in learning (\**P* < 0.05, \*\**P* < 0.01, two-sided Permutation test). Error bar represents SEM.

The cortical effects of PPC-rl inhibition depended on learning stage and response classes. In sensorimotor cortices, activity was significantly suppressed in beginner and intermediate stages but not in experts (Fig. 5f,g). Suppression was selective for delay and delay-reward neurons (Fig. 5f,g and Extended Data Fig. 6e-h), the very populations that normally strengthen in activity and connectivity with learning (Fig. 2c,d and 4d-f). These results suggest that PPC-rl boosts delay-related signals in sensorimotor circuits during association acquisition, a contribution that fades in experts.

To quantify how PPC-rl coordinates sensorimotor activity, we applied canonical correlation analysis (CCA) to analyze trial-by-trial co-fluctuations between PPC-rl and each sensorimotor region (Methods, Fig. 5h). We focused on the first CCA mode as it is the only significant mode that exhibits much higher canonical correlation than trial-shuffled datasets (P < 0.01, one-sided permutation test, Methods). Optogenetic inhibition of PPC-rl during the delay epoch markedly reduced the canonical correlation coefficient, with stronger effects for sensorimotor than non-sensorimotor cortices (Extended Data Fig. 7a,b). And this regional effect depended on learning stage, with robust effect in beginner and intermediate mice but absent in experts (P < 0.001, two-sided permutation test; Extended Data Fig. 7b). PPC-rl inhibition substantially attenuated co-fluctuation strength between PPC-rl and sensorimotor areas (Fig. 5h and Extended Data Fig. 7b), suggesting that PPC-rl is required to sustain coordinated trial-by-trial fluctuations with sensorimotor areas during learning.

We next examined intra– and inter-area co-fluctuations across somatosensory and motor cortices, and identified learning-stage specific reorganization (Fig. 5i,j). In beginner mice, PPC-rl activity aligned intra-somatosensory co-fluctuations (SSp-bfd↔SSp-others). In intermediates, this somatosensory intra-area coupling weakened, while a robust motor intra-area co-fluctuation emerged (MOs↔MOp). Concurrently, inter-area coupling between somatosensory and motor cortices appeared (SSp-bfd↔MOs). By the expert stage, correlations were significantly reduced (P < 0.001 vs. intermediate mice, two-sided permutation test), paralleling PPC-rl’s waning behavioral impact and decoding contributions (Fig. 2, 3). PPC-rl inhibition significantly diminished both intra– and inter-sensorimotor correlations in beginner and intermediate mice (control vs inhibition, SSp-bfd and SSp-others P < 0.01; MOs-MOp P < 0.01; SSp-bfd and MOs P<0.05; Fig. 5i,j), with little effects in experts. To assess whether this effect was specific to the CCA mode linking PPC-rl delay neurons to sensorimotor delay-reward neurons, we examined CCA modes built from other source-target combinations, including PPC-rl delay neurons paired with delay neurons in other areas, and sources in other areas (e.g., RSP delay neurons). PPC-rl inhibition did not modulate these alternative pairings (Extended Data Fig. 7c-f), confirming the specific link between PPC-rl delay and sensorimotor delay-reward activity. Together, these results show that PPC-rl enhances trial-by-trial co-fluctuations between somatosensory and motor cortices during learning. By aligning delay-related activity across areas, PPC-rl establishes temporally coincident sensorimotor activity that may facilitate Hebbian mechanisms linking sensory evidence to delayed action.

## PPC-rl persistence, PPC-rl dependent sensorimotor subspace communication and reward-gated Hebbian plasticity synergize to support delayed association

Our experimental results indicate that during learning, PPC-rl enhances trial-by-trial co-fluctuations between somatosensory and motor cortices, aligning their activity within a low-dimensional subspace. Because synaptic plasticity depends on temporally coincident activity^10–12^, such PPC-rl–mediated alignment could enable Hebbian strengthening between sensory and motor representations despite the sensory-motor delay. We therefore asked whether persistent PPC-rl activity, subspace alignment, and reward-gated synaptic updates are jointly sufficient to account for delayed sensorimotor association.

To test this, we constructed a minimal three-population recurrent network comprising SS (somatosensory, SSp-bfd and SSs), PPC-rl (association) and MO (motor, MOp and MOs) (Fig. 6 and Extended Data Fig. 8a-c, Methods). The architecture implements established anatomical and functional properties of the cortex: 1) whisker input drives SSp-bfd, 2) MO population activity controls licking output, 3) each area has local recurrent excitation, 4) long-range inter-areal projections are excitatory, 5) SSp-bfd and PPC-rl are reciprocally connected (Extended Data Fig. 9a-c)^35–38^, 6) PPC-rl and motor cortex are reciprocally connected (Extended Data Fig. 9a-c)^37–39^, and 7) PPC-rl has stronger recurrence to support stimulus-evoked persistent delay activity (Fig. 6a). Learning was implemented with a three-factor, reward-gated Hebbian rule^8,13^. Pre– and postsynaptic coactivation generates an eligibility trace that decays over time, and synaptic modification occurs only upon reward delivery, proportional to the reward signal and trace magnitude. In this framework, PPC-rl dependent persistence and sensorimotor subspace alignment supply the co-activity needed to accumulate eligibility traces, whereas reward converts those traces into lasting synaptic strengthening (Methods).

**Fig. 6.**
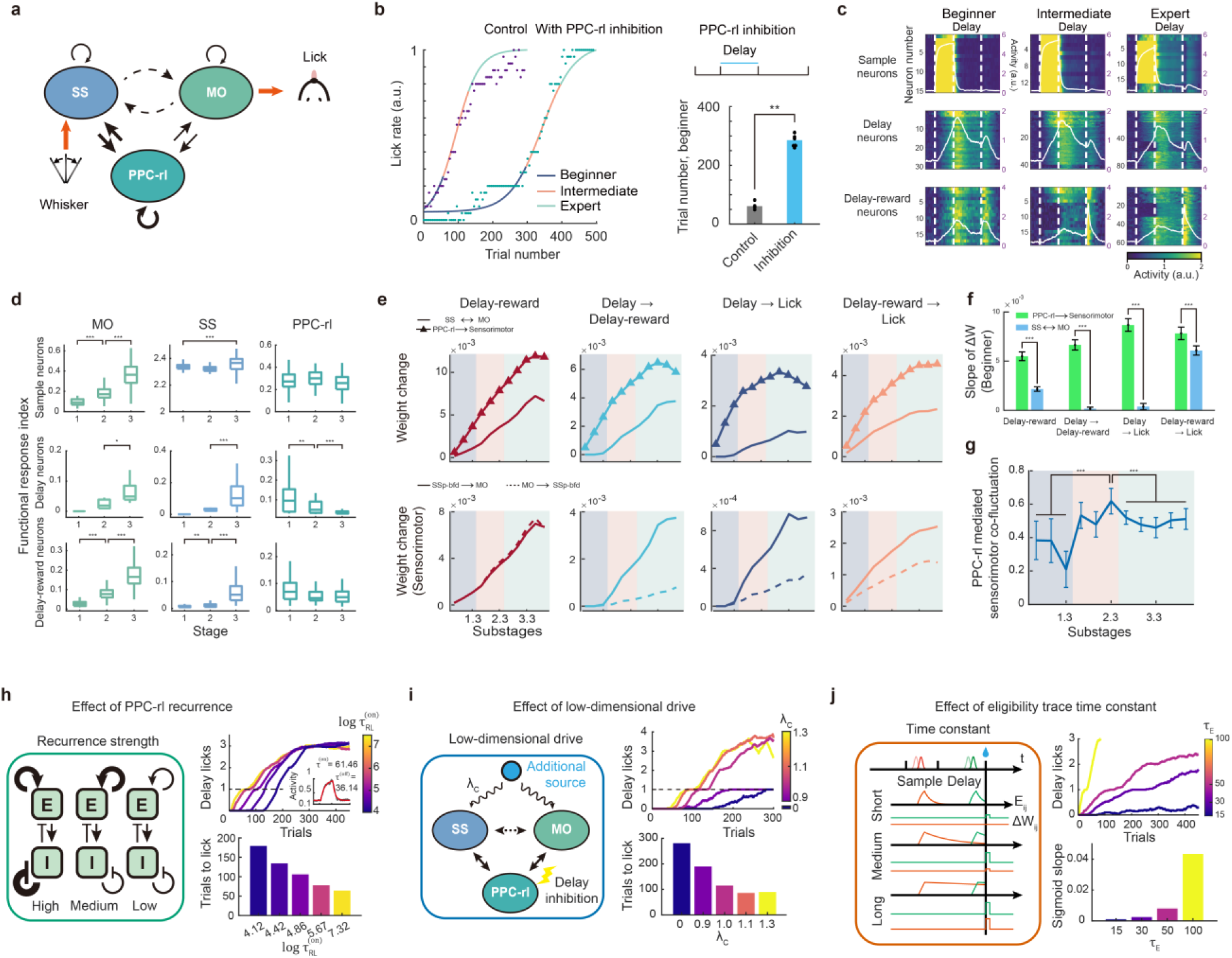
| PPC-rl persistence, PPC-rl dependent sensorimotor subspace communication and reward-gated Hebbian plasticity synergize to support delayed association. **a.** Schematic of three-area network. Each area (SS, PPC-rl, MO) is modeled as a recurrent rate network. Whisker input drives SS. MO activity controls licking output. Inter-areal projections are excitatory. PPC-rl has stronger recurrence to sustain delay activity. See methods for model details. **b.** Learning performance in control versus PPC-rl inhibition. Inhibition markedly prolongs the beginner stage and slows acquisition (\*\**P* < 0.01, two-sided Wilcoxon rank sum test). For all control and inhibition simulations, n = 6. **c.** Single-neuron activity across learning. Heatmaps show activity of sample, delay, and delay-reward selective units. Vertical lines separate behavioral epochs. White traces, mean for each class. Y-axis numbers, unit counts. **d.** Response amplitudes across areas and learning stages. Box and whisker plots show median (line), interquartile range (box), range (whiskers), and outliers (points). Significance is determined by two-sided Wilcoxon rank sum test (\**P* < 0.05, \*\**P* < 0.01, \*\*\**P* < 0.001). **e.** Connectivity strength increases with learning. Columns, connectivity for delay-reward, delay→delay-reward, delay→lick, and delay-reward→lick groups. Top rows, weight increase for intra-sensorimotor (SS↔MO, solid line) and PPC-rl→sensorimotor (triangle) connections. Bottom, intra-sensorimotor connections (SS→MO, solid; MO→SS, dashed). PPC-rl→sensorimotor strengthens earlier than SS↔MO connections. **f.** Weight increases faster and earlier for PPC-rl → sensorimotor compared with intra-sensorimotor connections (\*\*\**P* < 0.001, Wilcoxon rank sum test). **g.** Sensorimotor co-fluctuation strength across learning stages. Pearson correlation between SSp-bfd and MO population activity increases during learning and declines in experts. Sensorimotor co-fluctuation in substage 2.3 is significantly greater than that in stage 1 and stage 3 (\*\*\**P* < 0.001, Wilcoxon rank sum test). **h.** PPC-rl persistence dependence. Left, persistence modulation by changing strength of recurrence. Right, learning trajectories (top) and learning speed (bottom) for different persistence with different network time constants (see Methods). **i.** Rescue of PPC-rl inhibition by adding a low-dimensional sensorimotor drive. During inhibition, injecting a delay-specific low-dimensional drive into SS and MO restores learning. Left, schematic. Right, learning trajectories (top) and learning speed (bottom) for different drive strengths. **j.** Eligibility-trace dependence. Left, schematic of long, medium and short trace constants. Right, learning trajectories (top) and learning speed (bottom) for different trace durations.

With these biologically motivated elements in place, the simulation recapitulates major experimental observations across learning: 1) preparatory licking increases sigmoidally across training (Fig. 6b and Extended Data Fig. 8d), 2) SS and MO increases the delay and delay-reward activity (Fig. 6c,d and Extended Data Fig. 8e), 3) PPC-rl reduces delay persistence with learning (Fig. 6c,d and Extended Data Fig. 8e), which results from the strengthening of sensorimotor connections and synaptic normalization required to stabilize Hebbian growth, 4) PPC-rl inhibition slows learning by prolonging the beginner stage (Fig. 6b and Extended Data Fig. 8d), 5) synaptic strengthening first emerged in PPC→sensorimotor projections and only later within intra-sensorimotor connections for delay→delay-reward, delay→lick, and delay-reward→lick chains (Fig. 6e), 6) learning amplifies PPC-rl dependent sensory-motor co-fluctuations that declines in experts (Fig. 6f-g). Thus, the model captures both the emergence and eventual attenuation of PPC-rl’s functional influence observed experimentally.

Targeted model perturbations can further dissect the contribution of each component. We first examined the role of PPC-rl persistence. Increasing the persistence of PPC-rl delay activity accelerated learning (Fig. 6h and Extended Data Fig. 8f-g), whereas simulated PPC-rl inhibition prolonged the beginner stage (Fig. 6b). We next tested whether PPC-rl’s function could be attributed to its role in organizing low-dimensional subspace communication between somatosensory and motor populations. Although PPC-rl inhibition slowed learning, this deficit could be rescued by directly injecting a low-dimensional drive into SS and MO that imposed aligned activity during the delay epoch (Fig. 6i and Extended Data Fig. 8h,i). We finally modulated the plasticity rule by shortening the eligibility trace. Reducing the trace time constant markedly diminished, and in some cases abolished, the learning benefit of PPC-rl delay activity (Fig. 6j and Extended Data Fig. 8j,k).

Together, these model results reveal a division of labor during delayed sensorimotor learning: PPC-rl persistence sustains delay activity across the temporal gap, low-dimensional alignment channels this activity into sensorimotor co-fluctuations, and eligibility traces convert aligned co-fluctuations into lasting synaptic change at reward. Their interaction is jointly sufficient to reproduce the stage-dependent dynamics and connectivity reorganization observed *in vivo*.

## Model predictions and experimental verification

The model further generates two nontrivial predictions. First, although beginner mice exhibit little overt preparatory licking during the delay epoch, the model predicts that delay-epoch motor activity is nonetheless causally required for learning. This is because eligibility traces accumulate from temporally aligned sensorimotor co-activity primarily during the delay epoch preceding reward (Fig. 6). In simulations, inhibiting MO during the delay epoch markedly slowed acquisition (Fig. 7a-d, Methods), indicating that motor delay activity is not merely a readout of learning but a necessary substrate for building delayed association. Second, because PPC-rl provides a low-dimensional delay-specific input that aligns somatosensory and motor activity (Fig. 5), enhancing PPC-rl delay activity should accelerate learning by shortening early learning stages. Given the heterogeneity of PPC-rl response types, it is not obvious a priori that non-specific elevation of PPC-rl activity would facilitate acquisition. However, simulations demonstrated that enhancing PPC-rl delay-activity significantly reduced the duration of learning stages, thereby accelerating learning (Fig. 7e-h).

**Fig. 7.**
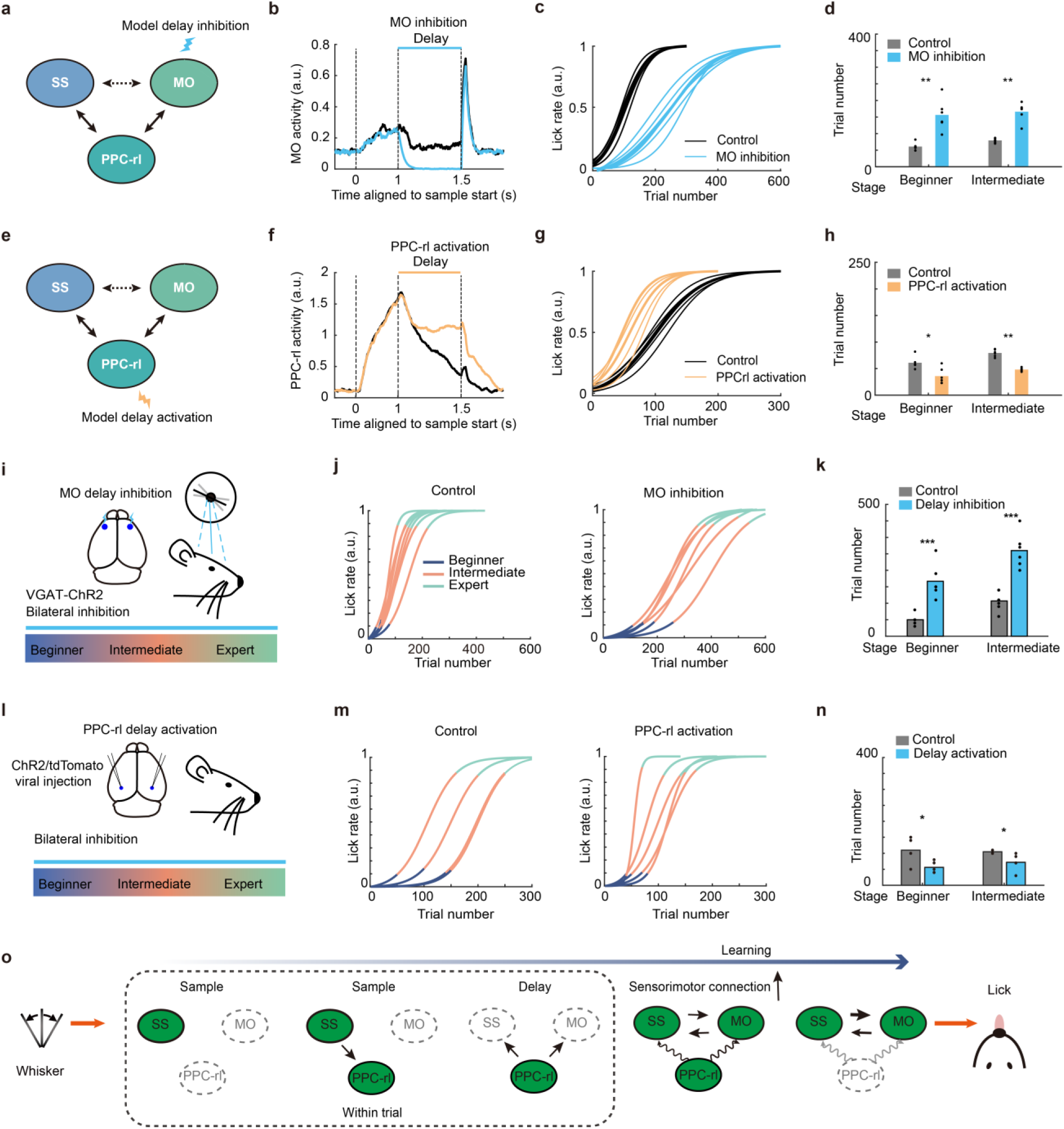
| Model simulation of MO inhibition and PPC-rl activation, and experimental validation. **a.** Simulation of MO inhibition during the delay epoch. Inhibition was applied on 75% of trials. Related to **b-d**. **b.** MO activity during control and inhibition. Thick lines, mean. For all control and model simulations, n = 6. Note that some lines or dots (in **d** and **h**) are overlapping due to small variability. **c.** Delay-epoch inhibition of MO slows learning. **d.** Quantification of the number of trials in the beginner and intermediate learning stages. MO delay-epoch inhibition prolonged duration of learning (\*\**P* < 0.01, two-sided Wilcoxon rank sum test). Same data as in **c**. **e.** Simulation of PPC-rl delay-epoch activation (100% of trials). Related to **f-h**. **f.** PPC-rl activity during control and activation. **g.** Delay-epoch activation of PPC-rl accelerates learning. **h.** Quantification of the number of trials in the beginner and intermediate learning stages. PPC-rl delay-epoch activation reduced the duration of learning (\**P* < 0.05, \*\**P* < 0.01, two-sided Wilcoxon rank sum test). Same data as in **g**. **i.** Schematic of optogenetic inhibition of MO during the delay epoch. Related to **j-k**. **j.** Experimental test of MO delay-epoch inhibition. Left, control mice without inhibition (n = 7 mice; note lines overlapping;). Right, inhibition during the delay epoch (n = 6 mice). Inhibition was applied on 75% of trials. Control mice and inhibition mice have the same genetic background. **k.** Quantification of the number of trials in each learning stage, demonstrating prolonged learning stages with MO delay-epoch inhibition (\*\*\**P* < 0.001, two-sided Wilcoxon rank sum test). **l.** Schematic of PPC-rl delay-epoch activation. Related to **m-n**. **m.** Experimental test of PPC-rl activation during the delay epoch. Left, control mice without stimulation (n = 4 mice; injected with tdTomato virus). Right, stimulation during the delay epoch (n = 5 mice; injected with ChR2 virus). Stimulation was applied on 100% of trials. **n.** Quantification of the number of trials in each learning stage. Delay-epoch activation of PPC-rl accelerates learning (\**P* < 0.05, two-sided Wilcoxon rank sum test). **o.** Mechanistic summary. Whisker input drives SS, which recruits PPC-rl. During the delay, PPC-rl sustains activity and aligns SS-MO dynamics within a communication subspace. This synchronized delay activity, gated by reward via an eligibility trace, strengthens SS-MO functional connections to link sensation to distal action. Green shading indicates activation.

We experimentally tested these predictions. Bilateral optogenetic inhibition of MO during the delay epoch significantly slowed learning (inhibition on 75% trials, with 25% intact trials used to quantify learning, Fig. 7i-k). This demonstrates that motor delay dynamics are causally required for strengthening sensorimotor associations rather than simply reflecting motor preparation. Conversely, optogenetic activation of PPC-rl during the delay epoch accelerated learning, leading to earlier emergence of preparatory licking and faster progression to expert performance (Fig. 7l-n). The shortening of early learning stage is not due to an acute increase of licking, as activation on probe trials did not evidently enhance preparatory licking (Extended Data Fig. 9e). Together, these results validate the core model predictions and emphasize the synergistic roles of PPC-rl persistence and motor delay dynamics in bridging the temporal gap during delayed sensorimotor learning.

## Discussion

Our study introduces a general strategy for dissecting distributed brain circuits of learning by combining activity-based large-scale screening, causal perturbation, and simultaneous readout during manipulation. First, cortex-wide imaging of >10,000 neurons revealed a distinctive learning-related reorganization in PPC-rl during a whisker-lick association with a temporal gap. Second, temporally precise optogenetic inhibition established that PPC-rl is required during acquisition but becomes dispensable for expert performance. Third, simultaneous inhibition and cortex-wide imaging showed that suppressing PPC-rl selectively attenuates delay-related sensorimotor activity in learners, revealing a stage-dependent mechanism. Fourth, analyses of trial-by-trial co-fluctuations and biologically grounded network modeling indicate that PPC-rl provides an instructive signal that strengthens somatosensory-motor coupling via reward-gated Hebbian-like mechanisms. Finally, the model generates mechanistic predictions that can be directly tested experimentally, enabling an iterative loop between theory and perturbation to refine circuit-level understanding. This framework, i.e. brain-wide recording to identify candidate hubs, targeted optogenetics to test causality, and concurrent perturbation with population recording to reveal circuit mechanisms, is readily compatible with approaches such as Neuropixels arrays that sample 1,000–2,000 neurons simultaneously at high temporal resolution^33,40–44^. Together, these advances provide a systematic path to map and test distributed computations across cortical and subcortical networks.

Conceptually, our results identify a circuit-level solution to the temporal credit assignment problem in sensorimotor learning. The “distal reward problem”^8^ or the “credit assignment problem” in the context of reinforcement learning^7^ can theoretically be supported by persistent or activity-silent working memory mechanisms, coordinated inter-areal communication subspaces, and synaptic eligibility traces gated by neuromodulators. Our data indicate that these mechanisms act synergistically. First, PPC-rl maintains a delay-period memory of the tactile stimulus (Fig. 7o). This signal depends on whisker-evoked SSp-bfd input, appears before the learned association, and is therefore not motor in origin. Because the somatosensory cortex alone does not sustain such persistent activity^44–47^, PPC-rl appears to transiently extend the temporal footprint of sensory input, bringing stimulus representations closer to reward for more effective learning. As learning progresses, mice initiate preparatory licking earlier, effectively reducing the temporal gap and the challenge of the distal reward problem. Indeed, selectively removing licking related eligibility traces in simulations significantly impaired learning (Extended Data Fig. 8l,m). Second, PPC-rl establishes a low-dimensional communication subspace that transiently aligns SS and MO population activity during the delay, increasing pre-post correlations when eligibility traces accumulate (Fig. 7o). This emerges from PPC-rl’s delay activity, that elevates both SS and MO delay-reward activity. Third, neuromodulatory outcome signals at reward convert accumulated eligibility into lasting synaptic change, consolidating sensorimotor pathways that renders PPC-rl dispensable for execution (Fig. 7o). There are direct connections between SSp and MOp, and parts of SSp and SSs also project to MOs^48,49^. After learning, axons of SSp/SSs have a higher efficiency in driving motor cortical neurons and in producing orofacial movements, suggesting strengthening of direct synapses (Extended Data Fig. 9f-o). The convergence of these elements is captured by our three-module recurrent network: PPC-rl persistence organizes cross-area co-fluctuations early in learning; reward-gated Hebbian plasticity strengthens SS→MO pathways; and as sensorimotor coupling consolidates, PPC-rl’s functional contribution regresses. The model framework is general; it can be generalized to architechtures incorporating intermediate structures (e.g. the prefrontal cortex or basal ganglia) between SS and MO (Extended Data Fig. 8n,o). Although alternative plasticity mechanisms such as behavioral time scale synaptic plasticity (BTSP) can drive rapid, one-shot changes over seconds^50^, the gradual, reward-dependent acquisition observed here is more consistent with incremental neocortical potentiation.

The posterior parietal cortex has been implicated in multisensory integration, attention, working memory, and decision-making^39,51–57^, yet causal studies have yielded dissociated outcomes. Inhibiting PPC can spare evidence accumulation and choice in some well-trained tasks^46,58–60^, but PPC is necessary for categorizing novel stimuli and contributes to trial-history dependent biases^61–63^. Our results identified a clear learning-specific role: PPC-rl is required during acquisition of a tactile sensorimotor association with a delay, but becomes dispensable once the behavior is well-learned. This pattern suggests that PPC scaffolds sensorimotor mapping by integrating sensory evidence and biasing downstream pathways until cortical sensorimotor circuits consolidate the association. Residual dependence on PPC likely scales with task demands: increasing task difficulty, the need for evidence accumulation, post-stimulus maintenance, or scaffolding for subspace communication engages PPC contribution even in well-trained animals^39,53,64,65^. Thus, PPC flexibly shifts from a teacher-like role during learning to a modulatory role during routine execution, with necessity determined by novelty, uncertainty, memory load as well as training duration.

How does PPC-rl drive the learning-related emergence of delay-reward activity in sensorimotor circuits? Anatomical tracing shows reciprocal connectivity between SSp-bfd/MOs and PPC-rl (Extended Data Fig. 9a-d)^35–38^. Additional polysynaptic routes can convey indirect influence of PPC-rl. First, PPC-rl projects to the posterior nucleus of the thalamus (PO), which projects broadly to motor cortex. Second, PPC-rl projects to the dorsolateral striatum, which might shape delay-reward activity in motor cortex via the output nuclei of the basal ganglia and thalamus. Through these pathways, PPC-rl may regulate inter-areal communication subspaces^18,66^, biasing correlated activity during the delay to promote selective strengthening of relevant pathways. Dissecting the relative contributions of direct corticocortical, thalamocortical, and striatal loops will be essential for linking PPC-rl persistence to consolidation of sensorimotor coupling.

Cortex-wide imaging has been instrumental for probing cortical dynamics during learning^22–24,67,68^. Previous widefield approaches, typically performed through intact translucent skull with dense neuronal labeling, capture population-averaged activity without single-cell resolution^21^. Sparse labeling combined with craniotomy glass window mitigates this limitation^19,69^. Here, we optimized sparsity by titrating the injected reagent dosage and validated cellular resolution against two-photon microscopy (Extended Data Fig. 2). Achieving single-cell resolution enables parsing activity into distinct functional neuron classes (sample, delay, delay–reward, lick) and tracking how their contributions evolve across learning, thereby revealing candidate hubs and stage-dependent reorganization that are typically obscured at the population level. Given the simplicity of the widefield imaging system, the availability of transgenic mouse lines, and mature pipelines for single neuron extraction, this approach is well positioned for broad adoption. Moreover, combining cortex-wide single-cell imaging with targeted causal manipulation provides a scalable framework for linking distributed activity patterns to the circuit mechanisms that drive learning.

### Limitations of the study

Our study has a few limitations that highlight important avenues for future research. First, while wide-field imaging with single-cell resolution enables cortex-wide monitoring of neural dynamics across learning, it is best suited for supragranular neurons. Due to light scattering and depth-dependent signal degradation, reliable cortex-wide imaging of deeper populations, particularly layer 5 projection neurons, remains technically challenging. Future optical advances will be necessary to determine how deeper output pathways contribute to the distributed learning processes described here. Second, we employed a simple detection task to isolate the core learning challenge of linking a sensory cue to delayed action and reward. In more complex discrimination paradigms requiring distinct behavioral responses to differentiate stimuli, PPC-rl activity would likely need to encode stimulus-specific patterns. Consequently, the broad optogenetic activation of PPC-rl that accelerated learning in our study might not generalize to tasks requiring precise, differential population codes. Finally, while we established that PPC-rl provides a crucial persistent signal to bridge the temporal gap during early learning, persistent activity becomes widely distributed across the brain in well-trained animals. Although our model framework is flexible to incorporate some intermediate area (e.g. representing prefrontal cortex or basal ganglia) between sensory and motor areas, future work is needed to determine how the PPC, prefrontal cortex, and subcortical structures like the basal ganglia interact as a unified network to support these distributed learning processes.

### Materials availability

All the reagents are available on request to Zengcai Guo.

### Data and code availability

The codes generated during this study will be available at GitHub with the following link: https://github.com/zqy97/Guolab_sensorimotor_association.git. The article includes all data generated during this study and all the raw data will be available from the corresponding author upon request. Any additional information required to reanalyze the data reported in this paper is available from the corresponding author upon request.

## Acknowledgements

We thank Xinyu Zhao and Yang Zhou for comments on the manuscript. This work was funded by Brain Science and Brain-like Intelligence Technology-National Science and Technology Major Project (2021ZD0203600); the National Natural Science Foundation of China (32525032, 32300852 and 32170998); and the support from THU-IDG/McGovern Open Laboratory of Shared Instruments for Brain Science.

## Author contributions

Conceptualization: ZVG and QZ; imaging experiments and data analyses: QZ; optogenetic experiments and data analyses: YW, JZ, AL, SL and QZ; electrophysiology and data analyses: YW; imaging with simultaneous optogenetic inhibition: QZ; model simulation: YH; anatomy: XY; funding acquisition, project administration and supervision: ZVG; writing: ZVG, QZ, YW and YH.

## Figures

**Video 1. Cortex-wide single-cell resolution imaging with simultaneous behavior monitoring during sensorimotor associative learning**.

This video illustrates multi-modal data collection and processing from a representative mouse in the expert learning stage. The top panel features the simultaneously captured facial video and the corresponding cortex-wide neuronal activity in the right hemisphere, along with the spatial map of all 16,000+ recorded neurons in Allen Mouse Common Coordinate Framework (CCFv3). The bottom panel displays the derived analyses, including a heatmap of real-time calcium activity for all neurons, the licking motion trajectory, and individual calcium traces from neurons identified as having the highest weights in the first principal component. This video is accelerated by 1.6x relative to real time speed.

## Methods

### Mice

All the experimental procedures were approved by the Use Committee and Institutional Animal Care at Tsinghua University, Beijing, China. Adult male mice at 8–16 postnatal weeks were housed in an animal facility (24 ℃ and 50% humidity) under a 12:12 reverse light:dark cycle and behaviorally tested during the dark phase. All behavioral experiments were performed on head-fixed, awake mice. For experiments of calcium imaging, and experiments with simultaneous imaging and optogenetic inhibition, we used 8 transgenic mice hybridized between Rasgrf2-2A-dCre (Jax#022864)^70^ mice and Ai148 (TIT2L-GC6f-ICL-tTA2)-D (JAX#030328)^71^ mice, that express Cre-dependent GCaMP6f genetically encoded calcium indicator. For experiments of optogenetic perturbation, we used 67 VGAT-ChR2-EYFP^72^ mice and 9 WT mice. For experiments of electrophysiology, we used 9 VGAT-ChR2-EYFP mice and 2 ReaChR mice.

### Surgery

Surgeries were conducted under 1–2% isoflurane anesthesia in accordance with institutional animal care guidelines. After surgery, flunixin meglumine (Sichuan Dingjian Animal Medicine Co., Ltd) was injected subcutaneously (1.25 mg/kg) during and after the surgery for at least three days to reduce inflammation^73^.

For electrophysiology and optogenetic inhibition, mice were prepared with a head bar and a clear-skull cap^46^. The scalp over the dorsal cortex was carefully removed to expose the skull, which was then cleaned and coated with a thin layer of cyanoacrylate adhesive (Krazy Glue, Elmer’s Products Inc) applied directly onto the intact bone surface. A custom-designed titanium head-bar was affixed approximately above the cerebellar region using dental acrylic (Jet Repair Acrylic, Lang Dental; Part# 1223-clear). To ensure efficient light transmission, the acrylic layer was polished and sealed with a thin coating of clear nail polish (Electron Microscopy Sciences, Part# 72180).

For experiments of cortex-wide imaging, a 8.0 x 8.0 mm^2^ (for full cortex-wide window) or 6.0 x 8.0 mm^2^ (for left hemisphere window) craniotomy was made with a skull drill. After removing the skull piece, a coverslip was implanted on the craniotomy region, and a titanium headpost was then cemented to the skull for head fixation. For bilateral PPC-rl inhibition with simultaneous cortex-wide imaging, we performed the craniotomy surgery in the right hemisphere and implanted an optical fiber above the left PPC-rl (AP 2.45 mm, ML 2.6, DV 0, Extended Data Fig. 6). This surgical procedure allowed delivery of light over the left and right PPC-rl for simultaneous bilateral inhibition. The location of PPC-rl was delineated beforehand based on intrinsic signal imaging (see detailed procedure in the section of Intrinsic signal imaging and brain registration).

### Behavior

After surgery recovery for about 3-5 days, mice underwent food and water restriction for 3-5 days (2.5-3 g food for slight food restriction and 1 ml water per day), and the body weight were kept at above 80% of the weight before restriction^74^. After ∼2 weeks following surgery, the cranial window became clear, and mice were ready for behavioural training and imaging. During training, mice would consume about 1.0 ml milk in a single training session, and would be feed extra food and water (1.5-2 g food and 0.5 ml water per day) to keep the body weight.

A stainless-steel pole (diameter, ∼0.9 mm) was positioned ∼5 mm lateral to the whisker pad at one side of the mice. Licking was monitored with an electric lickport that also delivered liquid reward (Fig. 1). The base of the metal pole was glued to the mirror of a galvanometer, and whisker stimulation was generated by the galvanometer-driven actuator that vibrated the pole. Before the training of associative learning, there were 3 shaping sessions where mice received the same whisker stimulation and milk delivery. In those shaping sessions, whisker stimulation and milk were delivered at random, uncorrelated times (i.e. no sensorimotor association).

The tactile-based sensorimotor association task comprised two trial types. On the detection trial, the pole was vibrated at 10 Hz for 1 second. The strength of stimulation was adjusted by changing the current input to the galvanometer (∼1800 °/s peak velocity) to achieve a vibration amplitude of ∼ 20 mm. After whisker stimulation, there was a 1.5 s delay epoch. At the end of delay epoch, a milk reward (∼5 µL) was delivered through the lickport. Mice consumed the milk reward in the reward epoch. We defined this period as the reward epoch (not response epoch) as mice developed preparatory licking and expert mice initiated licking in the sample and delay epochs. To prevent mice from predicting the presence of whisker stimulation, we used an inter trial interval (ITI) with a random duration in the range of 10 – 40 s. Catch trials used the same time structure but omitted both the whisker stimulation and milk reward. In each training session, there were about 150 detection trials, and same number of catch trials. Each session contained 10% reward omission trials. To prevent mice from using auditory cues associated with galvo mirror movement during optogenetic experiments, continuous white noise was played throughout the entire learning period. After mice reached the expert performance, they were further trained for one session to confirm performance stability. After completing training, 8 mice were tested under whisker-trimming conditions (Extended Data Fig. 1k,l).

Transgenic mice of Rasgrf2-2A-dCre x Ai148D were used for imaging (Fig. 1, 2, 4). Transgenic mice of VGAT-ChR2-EYFP were used for optogenetic inhibition (Fig. 3). We noted that the imaging mice learned slower compared with the inhibition control mice (without inhibition). Mice underwent more disturbance (e.g. when used for electrophysiological recording) typically performed less well compared with mice prepared in training rigs. For comparison of learning speed, we used mice with the same genetic background (e.g. VGAT-ChR2-EYFP with sham or optogenetic stimulation; wild type mice with ChR2 or tdTomato injection).

### Virus injection

To bilaterally inhibit PPC-rl, we injected rAAV-CaMKIIa-eNpHR3.0-mCherry-WPRE-pA (BrainVTA) to express Halorodopsin in excitatory neurons^75^. To bilaterally activate PPC-rl, we injected pAAV-CaMKIIa-hChR2(H134R)-EYFP (OBiO) to express Channelrhodopsin2 in excitatory neurons, and injected pAAV-hSyn-tdTomato (Taitool) in control mice. As PPC-rl is a small cortical area, we first performed intrinsic signal imaging to delineate the cortical boundaries of PPC-rl (see detailed procedure in the section of Intrinsic signal imaging and brain registration). The stereotaxic coordinates were around AP –2.45; ML 2.6; DV 0.4/0.8. For injections, glass pipettes (Drummond) were pulled and beveled to a sharp tip (outer diameter ∼ 20 – 30 μm). The pipette was back-filled with mineral oil and front-loaded with viral suspension immediately before injection. The total amount of 25 nl virus was injected per site with a rate ∼10 nl/min using a microinjector. After injection, the pipette remained in place for ∼5 min before slow withdrawal to avoid backflow.

To activate axon terminals projecting from SS to the MO, we injected rAAV-CaMKllα-Cre (Taitool) in the left SSp-bfd and SSs in ReaChR mice (JAX#026294). To target SSp-bfd and SSs, the left hemisphere was tilted down by 5° from the horizontal plane and the injection sites were AP –0.5 and –1.2; ML 3.0; DV 0.6 for SSp-bfd and AP –0.5 and –1.2; ML 4.0; DV 0.6/1.2 for SSs. The total amount of 100 nl virus was injected per site in SSp-bfd and 70 nl virus was injected per site in SSs.

### Histology

To confirm the cell type specificity of GCaMP expression in Rasgrf2-2A-dCre x Ai148 transgenic line, we performed immunostaining on mouse brain sections against parvalbumin (PV), somatostatin (SST), and vasoactive intestinal peptide (VIP). Mice were perfused transcardially with PBS followed by 4% PFA / 0.1 M PB. The brains were fixed overnight and sectioned on a microtome at 70 um thickness. The sections were then pre-incubated in 0.5% Triton X-100 in PB for 30 min, and blocked in 5% bovine serum albumin and 0.5% Triton X-100 in PB for 1 h at room temperature (∼22℃). Sections were then incubated overnight at 4℃ and two more hours at room temperature in 0.1% Triton X-100 containing the primary antibodies (PV: MAB1572, Sigma-Aldrich, SST: MAB354, Sigma-Aldrich, VIP: 20077, ImmunoStar). After washing with 0.01 mol/L PBS, sections were incubated for 2 h at room temperature in 0.1% Triton X-100 containing secondary antibodies. Images were acquired on a confocal microscope (FV3000, Olympus). GCaMP was rarely expressed in interneurons (∼0% in PV, ∼0% in SST, ∼6% in VIP, Extended Data Fig. 2b,c).

### Intrinsic signal imaging and brain registration

To align each brain to the Allen Mouse Common Coordinate Framework (CCFV3)^76^, we performed intrinsic imaging to map the primary somatosensory areas using the same setup for widefield imaging. Mice were under light anesthesia (1.2 % isoflurane). We first find the focus plane by visualizing surface vasculature under blue illumination (470 nm LED) and then switched to red illumination (660 nm, M660L4, Thorlabs) to measure activity-dependent reflectance through the craniotomy window. Images were acquired for 5 minutes at 10 Hz. We presented three types of tactile stimuli to the hemisphere contralateral to imaging: 1) the whisker stimulation with the same vibrating pole used in the behavioral task, 2) the forelimb stimulation, 3) the hindlimb stimulation of the paws (20 Hz for 5 s, 10-15 s between individual stimuli). For each stimulus, reflectance was averaged over a 5 s window after stimulus onset and compared to a 5 s baseline immediately preceding stimulus onset. These stimuli yielded three robust functional hotspots that served as anchor points (together with anatomical landmarks including the bregma, lamdba, and the midline) for accurate registration using geometric transformation. Registration employed only rigid (translation and rotation) and affine (scaling) transformations. Guided by the atlas borders and stereotaxic coordinates, we delineated 11 areas of interest including the primary motor cortex (MOp), secondary motor cortex (MOs), primary somatosensory cortex barrel field (SSp-bfd), other primary somatosensory cortex (SSp-others, including somatosensory nose, mouth, forelimb, hindlimb, and trunk), secondary somatosensory cortex (SSs), primary visual cortex (VISp), posterior parietal cortex anterior (PPC-a), posterior parietal cortex rostrolateral (PPC-rl), medial visual cortex (VISm, including visual cortex anterior medial and posterior medial), posteriolateral association cortex (PLA, including visual cortex anterior lateral and lateral), and retrosplenial cortex (RSP). After identifying these cortical areas, we applied the transformation parameters of image registration to get neuronal localization in CCF and cortical area identity.

### Extracellular electrophysiology

All recordings were made from head-fixed mice. Extracellular spikes were recorded by Neuropixels probes (Neuropixels 1.0, IMEC)^77^. The voltage signals were acquired through SpikeGLX (Bill Karsh and Tim Harris, Janelia Research Campus.).

VGAT-ChR2-EYFP mice were used for recording neural activity from the motor cortex (MO), SSp-bfd and PPC-rl. A small craniotomy (diameter, 2 mm) was made over the contralateral motor cortex (center, bregma AP 1.5, ML 2.5 mm to target the anterior lateral motor cortex ALM which is the premotor area for licking^46^) or the somatosensory cortex (center, bregma AP –1.7, ML –3 mm) one day prior to the recording session. In MO and SSp-bfd recording sessions, two NeuroPixels probes were inserted into the primary somatosensory cortex (SSp-bfd) and the motor cortex (MO) at an angle of 14° and to a depth of 1 mm, with both probes targeting the same hemisphere. Each recording session lasted for about 100 trials, each comprising a 1-second tactile stimulus (presented with 50% probability), followed by a random delay of 5– 20 seconds, and a reward delivered with 50% probability. For PPC-rl recording under SSp-bfd inhibition, the Neuropixels probe was inserted into the PPC-rl at an angle of 14° and to a depth of 1 mm. During recording, 50% trials were under SSp-bfd inhibition during the epoch with whisker stimulation. ReaChR mice were used for tagging neurons in MO which connected with SS. Recordings were performed around the left ALM while activating ReaChR-expressing axons from left SS. Photostimulation consisted of three powers (10, 20, 30 mW) and 5 light pulses (2-ms pulses with 200-ms interval). Photostimulation was performed in 60% of catch trials.

We used Kilosort3 to perform spike sorting^32^. Before feeding the data into Kilosort3, we first used CatGT (https://billkarsh.github.io/SpikeGLX/) to subtract the averaged value of the signals sampled at the same time to reduce the common noise. Kilosort3 could give many candidate clusters (even up to thousands). We used the pipeline developed by the Allen Institute for Brain Science (ephys_spike_sorting) to select high quality units: 1) removing repeated spikes in each cluster, 2) finding clusters that were likely to be noise, 3) calculating the properties of each cluster and 4) using the quality matrix to select units. Selected units need to meet the following criteria: 1) they were annotated by Kilosort3 as ‘Single Unit’, 2) they were not annotated as a noise cluster, 3) the average firing rate was higher than 1 Hz, 4) the average amplitude of the waveform was higher than 70 μV, 5) the estimated time of appearance was higher than 0.8.

We isolated 201 single-units from MO and 125 single-units from SSp-bfd across 16 behavior sessions in 5 mice. And we isolated 44 single-units from PPC-rl across 11 behavior sessions in 4 mice. Spike width was calculated as the trough-to-peak interval in the mean spike waveform. Neurons with a width <0.3 were regarded as putative fast spiking neurons. Neurons with a width >0.3 and <0.8 were regarded as putative pyramidal neurons.

For optical tagging experiments, tagged neurons were identified based on their responses to individual light pulses, following the established procedure in Yang et al^48^. For each pulse, spike counts were calculated in a 20 ms window before and after pulse onset. We focused the second to the fifth light pulses to reduce the potential photoartifacts associated with the initial light onset. The pre– and post-pulse spike counts were compared using a paired two-tailed t-test across trials. For neurons showing significant excitation, we further estimated the number of spikes evoked by each light pulse. Response latency was defined as the first time bin at which the baseline-subtracted firing rate reached 50% of its peak response. A neuron was classified as a tagged neuron if at least one pulse met all of the following criteria: the post-pulse spike count was significantly higher than the pre-pulse spike count (P < 0.01), the number of evoked spikes per pulse was greater than 0.2, and the response latency was shorter than 5 ms. We isolated 135 single-units with 12 tagged units across 8 sessions before learning and 102 single-units with 19 tagged units across 7 sessions after learning in 2 mice.

### Optogenetic experiments

For optogenetic inhibition experiments using VGAT-ChR2-EYFP mice, we delivered laser light through the clear-skull cap^46^. Blue light from a 473 nm laser (Chang Chun optics) was controlled by an acousto-optic modulator (AOM; MTS110-A3-VIS, Quanta Tech; extinction ratio 1:2,000) to produce a continuous square-wave stimulus with 1.5 mW power. A 2D scanning galvo system (GVS012, Thorlabs) was used to deliver light to the bilateral PPC-rl (AP –2.45, ML ±2.6 mm), PLA (AP –3.5, ML ±4 mm) or MO (AP 1.7/2.5, ML ±2.5/1.5 mm). Three protocols were used for inactivation. The first protocol was randomly inactivating 25% of the trials during the intermediate and expert stages. Same set of mice were used for testing the involvement during the intermediate and expert stages. Control trials are those without light inhibition. To test the effect of inhibition on learning speed, we used the second protocol that inactivated PPC-rl or PLA on every trial during the learning process. For control mice, which were also VGAT-ChR2-EYFP mice, a light-shielding cover was positioned above the animals’ head to shield the animal from laser perturbations. Different cohorts of mice are used for control, inhibition during the delay epoch, and inhibition during the sample and delay epochs (because mice only learned once). To test the effect of MO inhibition, we used the third protocol that inactivated MO on 75% of trials (100% of trials will make learning extremely slow). This provided 25% of control trials to estimate the learning stages.

In total, 5 mice were used for inhibition in beginner, intermediate and expert stages using for protocol 1. To examine the role of PPC-rl for learning with optogenetic inhibition using protocol 2, 25 mice were used, with 9 used for control experiments, 8 for inhibition during the delay epoch, and 8 for inhibition during the sample and delay epochs. In the no-delay task, PPC-rl was inhibited during the sample period. We used 9 mice for control experiments and 9 for inhibition. For inhibition of PLA, we used 6 optogenetic inhibition mice and 7 for control. For inhibition of MO, we used 6 optogenetic inhibition mice and the same cohort of control mice as for PLA inhibition.

To perform PPC-rl inhibition during cortex-wide imaging, we first performed intrinsic signal imaging to delineate the cortical locations of PPC-rl. We then injected rAAV-CaMKIIa-eNpHR3.0-mCherry-WPRE-pA (BrainVTA) bilaterally into PPC-rl. Laser light was used for bilateral inhibition (Aurora-300, Newtoon, 589 nm center wavelength that did not interfere imaging). The light beam was split into two paths using a 1×2 fiber optic splitter. The laser output from each fiber was ∼7 mW. We then connected one path to a 100 µm core optical fiber (NA = 0.22; Inper) above the left-side. Light from the other path was projected onto the right PPC-rl directly using a collimator. We randomly selected 40% trials to perform bilateral inhibition during the delay epoch (n = 3 mice).

The procedure PPC-rl activation was similar to PPC-rl inhibition during cortex-wide imaging. We bilaterally injected pAAV-CaMKIIa-hChR2(H134R)-EYFP (OBiO) or pAAV-hSyn-tdTomato (Taitool, for control mice) into PPC-rl. Stimulation was delivered using 488 nm laser light, split into two paths using a 1×2 fiber optic splitter to target both hemispheres simultaneously. Stimulation consisted of 10Hz 5ms pulse trains with an average laser power of 1.5 mW. We activated bilateral PPC-rl during the delay epoch of all trials throughout the learning process, and randomly selected 5% of these trials for testing the effect of PPC-rl activation alone (with whisker stimulation and reward omitted from those trials). Control mice underwent identical training and light delivery procedures. In total, we used 5 mice for PPC-rl activation and 4 for control.

### Cortex-wide imaging

We performed mesoscale cortex-wide imaging with simultaneous recording of mouse orofacial movements using behavioral cameras. A Teledyne FLIR Chameleon3 camera recorded mice facial movements and pupil size at 40 Hz. Neural activity was imaged using a macroscope (MVX10X, Olympus) equipped with a 1× planapochromatic objective (MVPLAPO 1X, Olympus) and a motorized focus unit (SZX2-FOA, Olympus). The macroscope was coupled to a scientific complementary metal–oxide–semiconductor (sCMOS) camera (Andor Zyla 4.2 Plus) with 2,048 × 2,048 pixels and a pixel size of 6.5µm x 6.5µm. Alternatively, we used a sCMOS camera from Hamamatsu (ORCA-Flash 4.0 V2) that has the same pixel count and pixel size. Illumination was provided by a 4-wavelength high-power LED Source (LED4D123, 405/470/565/625 nm center wavelength, Thorlabs) controlled by a LED driver (DC4104, Thorlabs). Blue light was filtered through a band pass excitation filter (BP460-480HQ, within the Olympus filter set) and was directed to a dichroic mirror (DM485HQ). The emitted fluorescence light was collected through an emission filter BA495-540HQ. The imaging system was configured with a 1.6× zoom and 1× objective to capture a ∼8.3 mm × 8.3 mm field of view (FOV), covering most of the dorsal cortex. At this magnification, each pixel corresponded to a spatial footprint of 4 × 4 µm², sufficient for cellular-resolution imaging. To minimize photodamage and photobleaching, we illuminated the cortex at 10 mW (significantly below the typical 50 mW used in comparable studies^69,78^). Images were acquired at 15 Hz to track GCaMP6f dynamics with sufficient temporal resolution.

We achieved single-cell resolution by inducing cortex-wide expression of GCaMP6f in L2/3 neurons using trimethoprim lactate salt (TMP), a drug that stabilizes the destabilized Cre recombinase^69^. Mice received oral gavage of TMP (0.1 mg per gram, using methylcellulose as a vehicle) once daily for three consecutive days. The dose was titrated to produce robust, consistent expression across the whole cortex. For imaging, the craniotomy window was leveled using a tip/tilt stage to ensure the entire FOV in the focal plane. Cortex-wide imaging was performed from the last shaping session through the last training session, with identical optical settings across sessions.

### Two photon validation

We used two-photon imaging to validate neuron positions extracted from widefield one-photon data. First, we acquired a 10-minute one-photon video to facilitate segmentation of putative neurons. We then performed two-photon imaging at three representative locations within the cranial window (rostral, middle, and caudal regions) using an Olympus FVMPE-RS microscope with a 10× objective (XLPLN10XSVMP, Olympus). For each site, we collected a z-stack of 24–28 planes at 10 μm steps. At each plane, 10 frames were averaged (1,024 × 1,024 pixels; 1.24 μm/pixel; 0.16 Hz effective frame rate) to increase the clarity of images. Excitation was set to 930 nm, and the emission light was filtered with a band-pass filter (BA495– 543). Neurons were segmented and counted manually using the standard-deviation projection of the one-photon video and the maximum-intensity projection of the two-photon z-stacks. One-photon images were registered to two-photon stacks via a linear geometric transform to maximize one-to-one correspondence between widefield ROIs and two-photon resolved somata. About 86% of widefield identified neurons could be resolved with two-photon imaging (Extended Data Fig. 2d-f). Notably, some well localized one-photon spots lacked clear two-photon counterparts. One possible explanation is that due to the slow speed of two-photon imaging of a 3D volume, the limited per-plane dwell time can miss sparsely active or transiently silent neurons. Thus, the 86% match rate is probably a lower bound, reinforcing that our approach achieves single-cell resolution.

### Behavioral data analysis

To delineate learning stages, we first calculated the rate of preparatory licking during the delay epoch. We then fitted a sigmoid function for the lick rate change during learning. Stage boundaries were determined based on the sigmoid function as follows. First, the midpoint of learning was defined as the trial of the maximum derivative (peak slope) where the learning improvement was the fastest. Second, two additional points were chosen equidistant from the midpoint. And the two points were located with their sigmoid values difference spanned 75% of the total sigmoid range (Extended Data Fig. 1e). Then the earlier point defined the boundary between stage 1 (beginner) and stage 2 (intermediate). And the later point was defined as the boundary between stage 2 and stage 3 (expert). This definition correlates well with the pattern of correlation matrix for lick rate. To obtain this matrix, lick-rate traces on each trial was smoothed using a sliding-window of ten consecutive trials across learning. Then pairwise Pearson correlations were calculated between averaged licking traces to generate a correlation matrix. The heatmap of the correlation matrix summarize similarity of licking patterns over the course of learning, revealing similar learning stages as those based on sigmoid fitting. To analyze the internal changes within each of the three states, we further subdivided each learning stage into three equal substages (stage 1.1 to stage 3.3).

For detection trials, a Hit was defined when at least one report lick occurred during the delay epoch. If mice did not lick throughout the delay epoch, the trial was determined to be a Miss. For catch trials, false alarm (FA) trials were defined as those where mice licked during the delay, whereas trials with no licks were defined as Correct Rejections (CR). Performance was quantified as [number of Hits + number of CRs] / 20 within a sliding window of 20 trials. The sensitivity was quantified as d-prime (d’) using the standard signal detection formulation based on Hit and FA rates.

### Calcium trace extraction and cell registration

To extract neural traces, we first aligned images across all recording sessions using image cropping and linear geometric transformation (based on control points of blood vessels). A typical session comprised ∼60,000-80,000 frames. To accommodate memory constraints by a workstation, we split an image stack into 16 patches that were processed in parallel and later stitched. The extraction pipeline was implemented in CaImAn, an open source tool to perform motion correction, cell segmentation, calcium activity extraction and spike activity deconvolution^79^. We applied linear motion correction, used the constrained non-negative matrix factorization (CNMF-E) to obtain spatial footprints and denoised ΔF/F traces, and deconvolved putative spike trains using OASIS (online active set method to infer spikes). Quality control removed components with low SNR, or excessive overlap with vasculature by applying a vascular mask. We also removed spots with low spatial correlation with spots identified from the high quality image of pixelwise standard deviation. After extracting data from multiple sessions, we registered cells with CellReg based on centroid positions and spatial footprint correlations^80^. We only kept matches exceeding both distance and correlation thresholds. For each animal, parameters (e.g., patch size/overlap, CNMF-E initialization K, gSig/gSiz, merge thresholds, and OASIS penalty) were similar across sessions to ensure comparability. Neural traces of registered cells were concatenated across sessions to yield a unified neuron-by-time matrix for downstream analyses (Extended Data Fig. 1m).

### Classification of functional response types

Neurons were classified into functional groups within each learning sub-stage (1.1, 1.2, 1.3, 2.1, 2.2, 2.3, 3.1, 3.2, 3.3). Within any 10-trial window of a sub-stage, a neuron was identified as lick-related if meeting the following two requirements. First, the Pearson correlation between the neuron’s ITI spike activity and lick rate exceeded r > 0.1 (P < 0.01, t-test). Second, the lick-locked response amplitude exceeded 4 standard deviations of its baseline. To distinguish neurons that putatively drive licking (“pre-lick”) from those activated by licking (“lick-evoked”), we computed the cross-correlation between ITI spike activity and lick rate. Neurons exhibiting a peak correlation with activity preceding licking were defined as pre-lick neurons (or lick neuron for subsequent analysis), whereas the rest was classified as lick-evoked. Consistently, activity of lick neurons was concentrated on lick-related motor areas (such as the anterior lateral motor cortex and tongue-jaw primary motor cortex) and the oral-facial primary somatosensory cortex (Fig. 2c). The proportions of these two populations remained stable throughout learning (Extended Data Fig. 3b). Lick-related neurons were excluded from other functional categories. For the remaining neurons, we tested epoch-specific responses in sample, delay, and reward periods. We averaged activity within each epoch and compared it to a matched pre-sample baseline. Within each 10-trial window, significance was assessed with paired Wilcoxon signed-rank tests (one-sided, P < 0.01). Neurons with decreased activity were labeled suppressed, whereas neurons were labeled activated if the average activity trace within the epoch exceeded 4 s.d. of the baseline. If multiple epochs met criteria, the neuron was marked with multi-epoch response (e.g., sample-delay, sample-reward, delay-reward, sample-delay-reward). We observed no sample-reward or sample-delay-reward neurons and only rare sample–delay neurons (Extended Data Fig. 3b). To quantify the dynamic change of functional groups, we calculated two properties for each substage: 1) the fraction of neurons whose response type matched their stage 3.3 label and 2) the fraction transitioning to the response type of the subsequent sub-stage (Extended Data Fig. 3e). For the experiment of PPC-rl inhibition during cortex-wide imaging, we classified response types based on control trials during individual substages.

After assigning response types, we mapped their spatial distributions across learning substages. For each substage and response class, we rendered a heatmap in which each neuron’s color reflected its baseline-subtracted mean activity within the class-specific epoch: sample neurons in the sample epoch, delay neurons in the delay epoch, delay–reward neurons in the reward epoch, reward neurons in the reward epoch, and lick-related neurons in the reward epoch. Because individual somata span only a few pixels, maps were smoothed with a Gaussian kernel (σ = 10 pixel, 265µm in CCF) to improve visibility. To quantify response strength of each response class, we defined a Functional Response Index (FRI) per neuron as FRI = A × P, where A is the neuron’s mean activity in its task-specific epoch (baseline-subtracted) and P is the fraction of neurons of the same response type within the corresponding brain region.

### Decoding of trial types

We used a linear support vector machine (SVM) to quantify how well task-responsive neurons in each brain area distinguished detection from catch trials across time within each learning stage (Fig. 2)^73^. For each time point (from −1.0 to +4.5 s relative to trial onset, sampled at the imaging frame rate) and for each area, decoding accuracy was estimated via 1,000-iteration bootstrap cross-validation. In each iteration, we randomly sampled 300 neurons pooled across animals (n = 5) for that area. The training set for the SVM was constructed using data from these neurons with 36 detection trials and 36 catch trials within a specific learning stage. The corresponding data from other 9 trials of each type served as the test set to evaluate the model’s prediction accuracy. The number of trials was determined by the smallest number of trials within that learning stage across animals. To quantify the classifier’s decoding accuracy for the sample, delay, and reward epochs, we averaged the decoding accuracy at each time point within that respective epoch. As a control, we computed the decoding accuracy using a dataset where neurons from all brain areas were pooled together. This process allowed us to compare and validate the specific encoding contribution of each individual brain area.

### Functional connectivity and communication subspace granger causality

To quantify the functional connection strength between neurons, we computed the noise correlation using their activity during the inter-trial interval (ITI). To avoid the influence of behavioral movements, we first calculated the motion energy of facial movement across learning process, and defined the ITI time windows in which the motion energy was below the mean ± one standard deviation as the resting-state. For each learning stage, the duration of resting-state was selected for FC computation. The functional connection strength for a given neuron pair was calculated as the Pearson correlation coefficient of their resting-state activity traces. The connection strength between different functional groups was calculated as the average of all pairwise connection strengths between the neurons belonging to the two groups. To further analyze changes in information flow between functional neuron populations during training, we computed a connectivity subspace Granger causality. To incorporate the strength of neuronal connections into the calculation, we first performed principal component analysis (PCA) on the functional connectivity matrix between the two populations. We selected the top PCA dimensions that collectively explained 40% of the cumulative variance and projected the activities of each neuronal population onto their respective principal components. Subsequently, we computed pairwise Granger causality between all pairs of the resulting one-dimensional activity traces from the two populations. Only significant pairs were kept for further calculation (P < 0.05, Granger test). The final connectivity subspace Granger causality value was obtained as the weighted average of all these pairwise causality values, with the weight for each pair being the product of the variances explained by the two respective principal components.

### Canonical correlations analysis

We used canonical correlation analysis (CCA) to quantify the functional co-fluctuation between sensorimotor areas based on a shared functional dimension defined by their relationship with PPC-rl delay neurons^19^. We also assessed the effect of optogenetic inhibition of PPC-rl on these co-fluctuations. The analysis proceeded as follows. First, for a given learning stage, we randomly selected three-quarters of the trials. We used CCA on the activity of PPC-rl delay neurons and the activity of delay-reward (or delay) neurons from a single sensorimotor area to identify the linear combinations with highest correlations. This defines the dimensions that best captures the co-fluctuation between PPC-rl and each of the sensorimotor areas. Next, the neural activities from the remaining one-quarter of trials (the test set) were projected onto individual CCA modes to calculate canonical correlation. To test the robustness of CCA modes, we compared the canonical correlation with that of trial-shuffled datasets. Only the canonical correlation for the first CCA mode was significantly higher than trial-shuffled datasets across learning stages (n = 141 – 360 trials, 15 – 569 cells per cortices in one learning stage, from 3 mice; P < 0.01 in stage 1, P < 0.001 in stage 2 and 3, one-sided permutation test). The first CCA mode also exhibited much higher canonical correlation than that of the following CCA modes (mode 2/mode 1, 37.9%; mode 3/mode1, 19.3%; Extended Data Fig. 7a). Thus, we focused on the first CCA mode for subsequent analysis. Next, we calculated the Pearson correlation coefficient between the projected test-set activities of one sensorimotor area and those of another sensorimotor area (projected to corresponding CCA mode). This correlation value represents the strength of internal co-fluctuation between the two sensorimotor areas within the functional subspace that is jointly correlated with the PPC-rl population. This entire procedure was repeated with 1,000 random bootstrap samples to obtain a robust estimate. To investigate the effect of optogenetic inhibition, we performed an analogous calculation using laser-on trials. In each bootstrap iteration, after projecting the control test-set data, we also randomly selected a matched number of laser-on trials. The activities from these laser-on trials for the same pair of sensorimotor areas were projected onto the same CCA mode (derived from control trials without optogenetic inhibition), and the Pearson correlation between their projections was computed. This allowed for a direct comparison of internal sensorimotor co-fluctuation strength between control and inhibition conditions. Inhibition of PPC-rl reduced canonical correlation coefficient, and this effect showed selectivity for sensorimotor areas compared with non-sensorimotor areas in learning (Extended Data Fig. 7b). To validate the specificity of our findings, we performed multiple control analyses. First, we computed the co-fluctuation between PPC-rl and brain regions outside the sensorimotor areas (Extended Data Fig. 7). Second, we repeated the entire CCA-based co-fluctuation analysis using the RSP as the source area instead of PPC-rl to assess its functional coupling with other brain regions, including sensorimotor areas. PPC-rl inhibition selectively reduced co-fluctuations between sensorimotor areas along CCA axes identified using PPC-rl delay activity and delay-reward activity in a sensorimotor target (Fig. 5 and Extended Data Fig. 7), with no comparable effects in other areas or along other CCA directions.

### Simulation of network dynamics

We constructed a rate model incorporating 3 inter-connected subnetworks, representing SSp-bfd, PPC-rl and MO (including both MOp and MOs), respectively (Fig. 6a, and Extended Data Fig. 8a-c). Each subnetwork contains *N_E_* excitatory (E) and *N_I_* inhibitory (I) neurons (*N_E_* = *N_I_*). The sample and delay periods in each simulation trial were set to be 200 and 300 time-steps respectively, in proportion to real epoch durations in experiments. Reward was delivered right after the delay offset that persisted for 20 time-steps, similar to reward delivery time (∼0.1s). For a single unit *i*, the dynamics of its firing rate is governed by

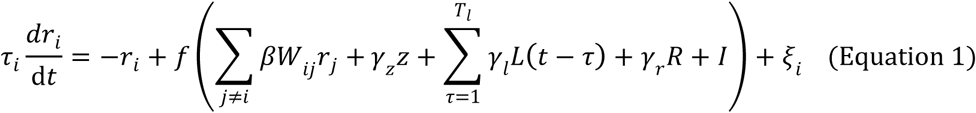

where the first term −*r_i_* reflects the exponential decay of its firing rate at the absence of any input, the second term is a transfer function *f*(⋅) with the argument summarizing inputs from other neurons, modulation effects from licking and reward neurons, and external input *I*(*t*). The third term denotes stochastic noise, with *σ_r_* a scaling factor and *ξ_i_*(*t*) following a zero-mean Ornstein-Uhlenbeck process, d*ξ_i_* = −*ξ_i_*d*t* + *σ_r_*d*W*, that evolves at each time step based on a Weiner process. For the second term, *W_ij_* are connection weights from unit *j* to unit *i* with elements non-negative and *W_ii_* = 0 to exclude self-connections. The connections originating from excitatory neurons and inhibitory neurons are specified by the value of *β*, with *β* > 0 for excitatory synapses and *β* < 0 for inhibitory synapses. The synapses from the same neuron have the same sign, respecting Dale’s law. To account for difference of inter-regional and local communication, we incorporated synaptic delay for all long-range excitatory connections, sampling from a discrete uniform distribution on [5,20]. Lick and reward modulate the network dynamics, and we explicitly simulated this process using *z*(*t*), *L*(*t*) and *R*(*t*), with *z*(*t*) the contribution of activity that drives lick (‘pre-lick’), *L*(*t*) lick-evoked activity, and *R*(*t*) as reward (Extended Data Fig. 8b). The modulation strength are constants *γ_z_*, *γ_l_* and *γ_r_*, and the duration of lick-triggered modulation is *T_l_*. The external sensory input *I*(*t*) was applied to a subpopulation of SSp-bfd. The nonlinear transfer function *f*(⋅) was set to be a rectified saturating sigmoid function with gain *α*_1_, offset *x*_0_ and slope *ρ*, thresholded at 0 (Equation 2),

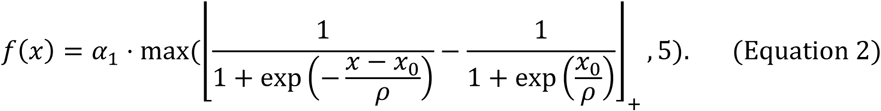

The connection weights among all local neuron pairs were initialized by a mean-shifted standard normal distribution (*μ*=0.2), clipped at 0.3. Therefore, local connections could exist in all types of neuron pairs. In contrast, only excitatory neurons were allowed for long range projection. The network was initialized with 15% “pre-lick”, 15% “lick-evoked” and 50% “reward” neurons (randomly chosen), similar to classes identified from calcium imaging data. Neurons in pre-lick group and lick-evoked group did not overlap.

Input to the network modulated half of the E neurons in SSp-bfd network, reflecting that only a fraction of neurons in the somatosensory cortex directly receive whisker information from the thalamus. By the same scheme, excitatory neurons in PPC-rl and MO were evenly split, with only one group receiving direct projections from SSp-bfd, whereas connection weights between SSp-bfd and indirect PPC-rl and MO groups were fixed to be 0. Neuron dynamics were transformed to licking outputs by introducing an artificial licking unit, whose activity *q*(*t*) was determined by a linear combination of all excitatory MO neurons, i.e., *q*(*t*) = *α*_2_ ⋅ *W_q_r*^(*E*^ ^from^ ^MO)^(*t*). Licking events were produced by a stochastic process, integrating *q* in each trial after stimulus offset in the presence of noise (a Weiner process with scaling factor *σ_z_*), formulated by

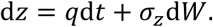

Once *z*(*t*) crossed a threshold *θ*_0_, a licking event would be generated, yielding *L*(*t*) = 1. A refractory period with duration *t_refr_* = 30 would be initiated upon lick onset, during which the licking integration was abolished.

In Extended Data Fig. 8n-o, we simulated a four-region model to evaluate the role of PPC-rl when SS and MO were not directly connected. To bridge the two regions, an additional subnetwork was introduced as a sensorimotor interface (SMI), making the SS–SMI–MO pathway conceptually analogous to the somatosensory-prefrontal cortex-motor cortex pathway, or the cortico-basal ganglia-thalamocortical pathway. Direct SS→MO connections (E→E and E→I) were removed, forcing all feedforward signals to traverse SMI. SMI contained the same number of E and I units as the other regions. The connection weights among SS, SMI and MO were initialized using the same clipped mean-shifted standard normal distribution, but with the following restrictions: 1) Feedback connection from SMI to SS was removed; 2) While in principle all I neurons in SMI could be modulated by SS, E neurons were separated into two partially overlapping subpopulations: an input group (75%) receiving SS afferents, and an output group (75%) projecting to MO, sharing 50% of total E units in SMI; 3) Local recurrent excitatory connections were abolished for SMI neurons.

### Synaptic plasticity

In our model, E-to-E and I-to-E connections were plastic, while E-to-I and I-to-I connections were fixed for simplicity (Extended Data Fig. 8c). Such implementation has been widely adopted by several modeling studies to explain basic neuronal properties including response flexibility and ensemble formation^81,82^.

E-E connections evolve on a trial-by-trial basis, following a variant of Hebbian rule. At each time-step, the degree of covariation between excitatory neuron *j* and *i* were estimated with a time window *T_e_* and added to an eligibility term *e_ij_*, whose dynamics is governed by

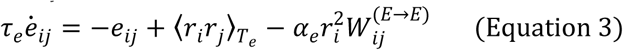

to produce the eligibility trace. The third term in the equation helps synaptic homeostasis and guarantees numerical stability. At reward offset, the eligibility trace was transformed into synaptic weight changes according to

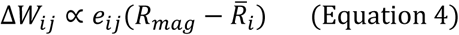

with learning rate *α_EE_*. *R_mag_* is a constant reward magnitude. R̅*_i_* is the estimated reward proportional to the average late delay activity of post-synaptic neuron *i*, calculated from 150 time-steps prior to reward onset. Meanwhile, dynamics of I-to-E synapses were slightly different. For example, the weight change from inhibitory neuron *j* to excitatory neuron *i* was computed by classical Hebbian rule, i.e., 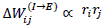, with learning rate *α_IE_*, updating at every time-step.

### Learning stage definition for in silico model

The effect of learning could be read out by the number of lick events in the delay period (*L_d_*). Across training, the trajectory of *L_d_* was fit by a four-parameter sigmoid function

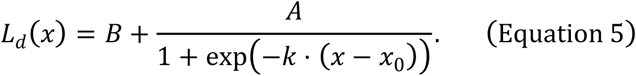

Here, *A* sets the maximum lick count increment in expert animals, *k* determines the growth rate, and was referred to “sigmoid slope” hereafter. *B* is the bias term, often greater than zero as *L_d_* rose to 1 in only a few trials in normal conditions after simulation start. Subsequently, three learning stages were defined using the same criteria as those applied to mice behavioral data (see Methods, Behavioral data analysis).

### Determination of response types for artificial units

In each trial, network units were classified based on their response profile (Fig. 6c,d and Extended Data Fig. 8e). Note that the pre-defined “pre-lick”, “lick-evoked” and “reward” groups were excluded here. First, a neuron was considered as task-activated if its average activity in sample, delay or early reward epoch (100 time-steps after reward onset) significantly exceeded baseline level (average activity in 100 time-steps before stimulus onset). Significance was determined by a 3-sigma rule. For example, if average sample and baseline activity for neuron *i* was denoted 〈*r_i_*〉*_S_*, 〈*r_i_* 〉*_B_*, neuron *i* was regarded as active in sample period if 〈*r_i_*〉*_s_*-〈*r_i_*〉*_B_* was greater than 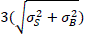, where 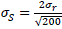 and 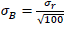 were the corresponding noise level. Next, peak time for every task-activated neuron’s activity, prior to reward onset, was quantified. Neurons were labeled as “sample-active” population if their activities peaked in sample period, and average activities in sample period were significantly greater than those in delay period, also decided by the 3-sigma rule. Likewise, neurons were labeled as “delay-active” population if their activities were prominent in delay epoch, compared to sample epoch. Furthermore, the “delay-reward” population was distinguished from other “delay-active” neurons due to their activation in early reward epoch, significantly greater than the average late delay activity. The response types defined here served as a basis for later analysis, such as quantification of learning related inter-regional connection changes and PPC-mediated co-fluctuation analysis (Fig. 6e-g).

To reveal the learning related changes of neural response strength of all three types of neurons, we next calculated the Functional Response Index for each region, defined as the average firing rates of all neurons in a specific population within a corresponding time window times the proportion of this population (the full sample period for “sample-active” and the full delay period for “delay-reward” population, same as Fig. 2d; the first 100 time-steps of delay period for “delay-active”, and changing this to the full delay period produces similar results).

### Learning related changes in connectivity between functional populations in simulation

Different from the analysis for experimental data, the connectivity between neuron pairs could be directly retrieved by the weight matrix. We focused our analysis on E-E connections. To summarize the learning related dynamics of connection weights, we first divided three learning stages to finer substages (3 substages for stage 1 and 2, 5 substages for stage 3). More substages were assigned to Stage 3 to balance trial counts per substage. Next, in each substage, we block-partitioned the connectivity matrix by response type and regional labels. The average value of all blocked matrix was then computed as the connection strength from some “sender” population to “receiver” population. We focused on analysis of “delay-reward to delay-reward”, “delay to delay-reward”, “delay to pre-lick” and “delay-reward to pre-lick” pairs as their connectivity reflect the internal information flow that guided sensorimotor transformation. Changes of connection strength between typical populations over training were displayed after subtracting the initial value and smoothed with a window length of 3 for visualization (Fig. 6e).

### Analysis of PPC-rl mediated communication subspace between SS and MO neurons in simulation

Within each learning stage, we randomly sampled 75% of trials as a training set. CCA was then applied to the activity of PPC-rl “delay-active” neurons and to the activity of “delay-reward” neurons from either SS or MO area, obtaining the first canonical variates that defines a one-dimensional axis that best captures co-fluctuation between PPC-rl and the corresponding sensorimotor area. We then projected the neural activity from the held-out 25% of trials (test set) for two different sensorimotor areas onto this CCA axis, and computed the Pearson correlation between the projected time series for the two areas (Fig. 6g).

### Effect of key model parameters on learning process

We examined three key factors that can be tuned to shape learning: the eligibility time constant, PPC-rl network recurrence, and the strength of low-dimensional PPC-rl mediated coordination between SS and MO (Fig. 6h-j). The working hypothesis is that PPC-rl recurrence generates delay-epoch persistence, which synchronizes SSp-bfd and MO activity, increases their coactivation, and creates eligibility traces that decay until reward delivery.

Firstly, the PPC-rl network recurrence was tuned by adjusting the strength of E-E and I-I connections in PPC-rl (Fig. 6i). The E-E connection helps persistence of external excitation, while I-I connection delays inhibitory recruitment that could otherwise suppress E activity. To quantify network recurrence, we used a mean-field approach and fit an asymmetric first-order linear response to the PPC-rl population activity using the following model,

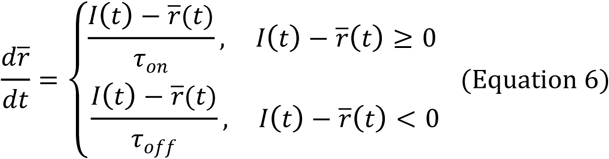

Here, *τ_on_* and *τ_off_* refer to time constants during the rising and falling phases following external input *I*(*t*). Model fitting was performed using MATLAB function lsqcurvefit. We reported *τ_on_* as the primary index of network recurrence, but qualitatively similar result was obtained using *τ_off_* (Extended Data Fig. 8f,g). To probe the impact on learning, we decreased the I-I coupling from 2.0 to 1.0 to 0.5. Then starting from the lowest I-I coupling, we reduced E-E coupling from 1.5 to 1.0 to 0.5, yielding 5 parameterizations with *τ_on_* = 61.46, 83.32, 128.82, 289.36 and 1512.71. Learning trajectories were computed as delay licks with a running average of 50 trials. Learning speed was quantified as the trial at which the trajectory reached 1 and maintained above. Note that, although the learning trajectory exceeded 1 faster under larger *τ_on_*, it ascended more gently to maximum. This effect should be attributed to the synaptic update rule used here (equation 4), as the appearance of lick would increase R̅*_i_* for many sensorimotor neurons, thus decreasing the third factor magnitude and the overall learning rate.

Secondly, to isolate the function of PPC-rl delay activity, we inhibited it by selectively activating PPC-rl inhibitory neurons during delay period, with an external current applied to all inhibitory neurons. This manipulation abolished early sensorimotor synchrony, and significantly postponed learning (Fig. 6b). To establish the causal contribution of early sensorimotor synchrony to learning performance, we introduced an additional low-dimensional, delay-specific drive to SSp-bfd and MO at each time-step of the delay period, while maintaining PPC-rl inhibition. We varied the strength of modulation (*λ_C_*) from 0.9 to 1.3, and assessed learning performance using the same criterion as above (Extended Data Fig. 8h). Interestingly, learning curves were qualitatively different for *λ_C_* < 1 and *λ_C_* ≥ 1, suggesting a critical synchrony threshold. Fitting these trajectories with the sigmoid function (Equation 5) revealed that *λ_C_* ≥ 1 yielded larger asymptotes with faster overall learning rates. In contrast, *λ_C_* < 1 yielded a slower approach to a ceiling of 1 (Extended Data Fig. 8i).

Lastly, we assessed how the time constant of eligibility trace (*τ_e_* in Equation 3) affected learning, by simulating the learning curve with *τ_e_* =15, 30, 50, 100 and 200 (Fig. 6j and Extended Data Fig. 8j,k). Conceptually, a small *τ_e_* restricts plasticity to neurons pairs coactive in the late delay period, whereas large *τ_e_* broadens the temporal credit-assignment window so that coactivity at any time during the trial epoch strengthens connections. In the first condition, training may be ineffective because potentiation is limited to coincident events close to reward and can concurrently potentiate I-to-E connections, yielding activation loss over the course of training. On the contrary, in the second condition, learning can quickly take place, making performance quickly reached the criteria, but the network will become overly excitable and tends to run away. Therefore, only *τ_e_* within certain critical range can support effective learning and stable performance. We quantified the effect of learning by fitting the sigmoid function (Equation 5) to the corresponding learning trajectory. We found performance could not reach the 1-lick criterion when *τ_e_* = 15, while the network activities ran away and produced epileptic behavior after 45 trials when *τ_e_* = 200. For remaining values, learning was effective and proceeded exponentially faster for larger *τ_e_*. In summary, only *τ_e_* within a bounded range supports robust and stable learning, aligning with our conceptual predictions.

### Network manipulation

To elucidate the functional contribution of MO, PPC-rl as well as preparatory licking in the model, we simulated four additional perturbation experiments (Exp 1, MO inhibition, Fig. 7a-d; Exp 2, PPC-rl activation, Fig. 7e-h; Exp 3, removing licking during delay, Extended Data Fig. 8l,m; Exp 4, PPC-rl inhibition using the four-area model, Extended Data Fig. 8n,o). In Exp1, MO inhibition was achieved by selectively activating MO inhibitory neurons on 75% of total trials. To simulate PPC-rl activation in Exp 2, we applied a weak external current to one fourth of randomly selected E neurons on every trial. The current amplitude was 20% of that used for inhibitory neurons in the PPC-rl inhibition simulation. This reduced amplitude was chosen to prevent runaway excitation and maintain activity within physiological bounds, ensuring that the manipulation enhanced delay activity without destabilizing network dynamics.

To model the effect of preparatory licking in Exp 3, we blocked lick-related eligibility trace, corresponding to setting the 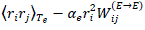 term in Equation 3 to be 0 for all incoming and outgoing synapses of lick-related neurons in delay epoch. Pre-lick neurons were modulated by the lick unit during preparatory period, and eligibility increment during time-steps where *z*(*t*) > 0 was set to zero. Lick-evoked neurons were directly modulated by licking events and the eligibility term did not increment for [0, *T_l_* + *T_e_*] after licking event. In Exp 4, PPC-rl inhibition was achieved by selectively activating PPC-rl inhibitory neurons during delay epoch. The strength of inhibitory modulation was identical to PPC-rl inhibition using the three-area model (Fig. 6a,b).

### Statistics

All the statistical tests were performed using Matlab R2021b (Mathworks). Statistics used for comparing behavioral performance was mentioned in corresponding places. Briefly, we used two-sided Wilcoxon sign rank test to compare within-stage trial-by-trial lick rate coefficients, cross-stage lick rate coefficients, and lick rates between control and optogenetic perturbation conditions, and used Kruskal-Wallis ANOVA test, followed by a post hoc Dunn’s multiple comparison test to compare performance and d’ in three learning stages.

To quantify functional response index, we used hierarchical random sampling (bootstrap) to account for mice and trial variability^46^. We first sampled the same number of animals with replacement, and then from each selected animal we randomly sampled trials with replacement. We repeated this process for 100,000 times but the results are consistent by reducing this number by 10-100 times. The P value for each condition was the fraction of times the response index during one learning stage was above or below a different learning stage (one-sided tests). Random sampling without considering mice and trial variability can render minimal difference significant, and thus this procedure presents a stringent test of significance. To compare the decoding performance of individual areas, we first compared individual cortices with the global population and then compared each significant area with the population of significant areas. We randomly sampled 300 neurons (without replacement) pooled across animals to account for neuron number difference in different areas. As the overlapped neurons between two random samplings was about 5%, and thus were treated as independent. We repeated this process for 100 times to avoid extensive usage of overlapped neurons.

In functional connectivity analysis, resting-state connection strength of 835,441,216 neuron pairs from 5 mice were calculated. Two-sided Wilcoxon sign rank test was used for screening significant connection among functional groups, and Kruskal-Wallis ANOVA test, followed by a post hoc Dunn’s multiple comparison was used to compare FC in sensorimotor cortices, FC between PPC-rl and sensorimotor areas and FC among other cortical regions. In Granger causality analysis, only significant pairs were kept for further calculation (P < 0.05, Granger test). Kruskal-Wallis ANOVA test, followed by a post hoc Dunn’s multiple comparison was used to compare FC and Granger causality in three learning stages between functional groups of interest.

In CCA analysis, we applied one sided permutation test to test the robustness of CCA modes as the previous study^19^, and applied two sided permutation test to compare difference between control trials and PPC-rl inhibition trials. Normally distributed data were presented as mean values together with the standard error of the mean. In experiments with optogenetic inhibition of PPC-rl during beginner, intermediate and expert stages (protocol 1), lick rate during the delay period was compared using two sided Wilcoxon rank sum test. For the optogenetic manipulation applied throughout the entire learning process (protocol 2), we quantified the trial number of each mouse at different learning stages. Statistical significance between light-on and control conditions was tested using two sided Wilcoxon rank sum test. No statistical methods were used to predetermine sample size, but our sample sizes (mice number) are similar to those reported in previous publications^19,26^. Generally, we used 3-5 mice for experiments where we can compare perturbation effects in the same animals (perturbation and control trials were interleaved), while we used 6-10 mice if we need to compare learning process in different animals (control vs perturbation).

For model simulations, we first quantified the trial number of each round of simulation, and used two-sided Wilcoxon rank-sum test to analyze the statistical significance of learning speed between control and perturbation experiments (n=6 were performed for all conditions because the outcomes of individual simulations were relatively consistent).

## Extended Data Figures

**Extended Data Fig. 1.**
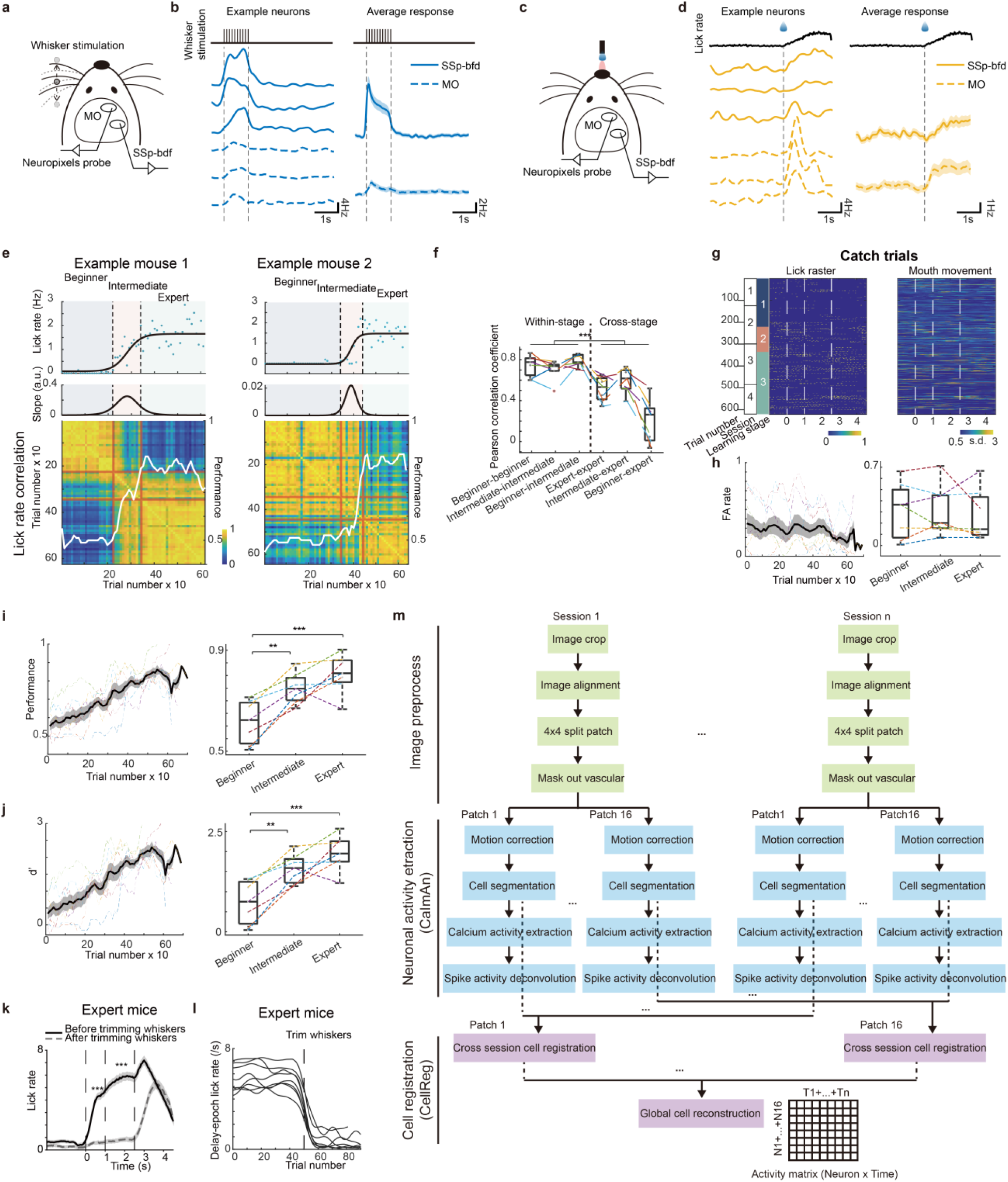
| Behavior characterization across learning stages. **a-d**. Tactile– and lick-modulated neurons rarely overlap in activity for naïve mice. **a**. Schematic of Neuropixels recording from somatosensory and motor cortex during whisker stimulation by a vibrating pole. **b**. Tactile-modulated neurons from the somatosensory barrel cortex (SSp-bfd) and motor cortex (MO). Left, example single neurons. Right, population average (n = 125 neurons in SSp-bfd; n = 201 in MO). Shading, standard error of mean (SEM). **c**. Schematic of recording during licking. **d**. Lick-modulated neurons. Same format as in **b**. **e**. Learning stage partition was concordant with performance measures. Top two rows: sigmoid fit of lick rate and its derivative (the left two panels reproduced from Fig. 1c). Dashed lines separates learning stages. The third row: heatmap showing trial-to-trial licking rate correlation. To calculate the correlation, we used blocks of 10 trials. Light red lines indicate the same trial indices as the dashed boundaries above. White traces are overlaid on each panel to show the performance of example mice. **f**. Pearson correlation coefficients for different learning stages. The within-stage coefficients are significantly higher than cross-stage coefficients (n = 7 mice; \*\*\**P* < 0.001; Wilcoxon rank sum test). Box plot shows the median, upper and lower quartiles. Whiskers extending from the box indicating the upper and lower data points. Dashed lines, individual mice. **g**. Behavior in catch trials across stages from the same mouse as in Fig. 1e. Left: lick raster (yellow ticks) for each trial. Right: z-scored mouth-movement traces. **h**. Left: false alarm rate across trials during associative learning. Dashed lines, individual mice. Solid line: mean. Shading: SEM. Right: false alarm rate in beginner, intermediate, and expert mice. (n = 7, *P* > 0.05; Kruskal-Wallis ANOVA test, followed by a post hoc Dunn’s multiple comparison test). Same box-plot format as in **f**. **i**. Left: learning curves across trials during associative learning. Dashed lines, individual mice. Solid line: mean. Shading: SEM. Right: behavioral performance in beginner, intermediate, and expert mice (n = 7, \*\**P* < 0.01; \*\*\**P* < 0.001; Kruskal-Wallis ANOVA test, followed by a post hoc Dunn’s multiple comparison test). Same box-plot format as in **f**. **j**. Same format and statistics as in **i**, but for performance characterized using d’. **k**. Lick rate before and after trimming whiskers for expert mice (n = 8, \*\*\**P* < 0.001, t-test). **l**. Change in preparatory lick rate after trimming whiskers for individual mice (n = 8). **m**. Pipeline of calcium activity extraction. First, we adopted image crop and linear geometric transformation to align images across all recorded sessions. After preprocessing raw fluorescence images, we used the constrained non-negative matrix factorization (CNMF) algorithm to extract neuronal activity traces. The extraction pipeline is implemented in CaImAn, an open source tool to perform motion correction, cell segmentation, calcium activity extraction and spike activity deconvolution (see Methods). After extracting data from multiple sessions, we used CellReg for cross-session cell registration (see Methods). Neural activity of registered cells was concatenated to form a single matrix.

**Extended Data Fig. 2.**
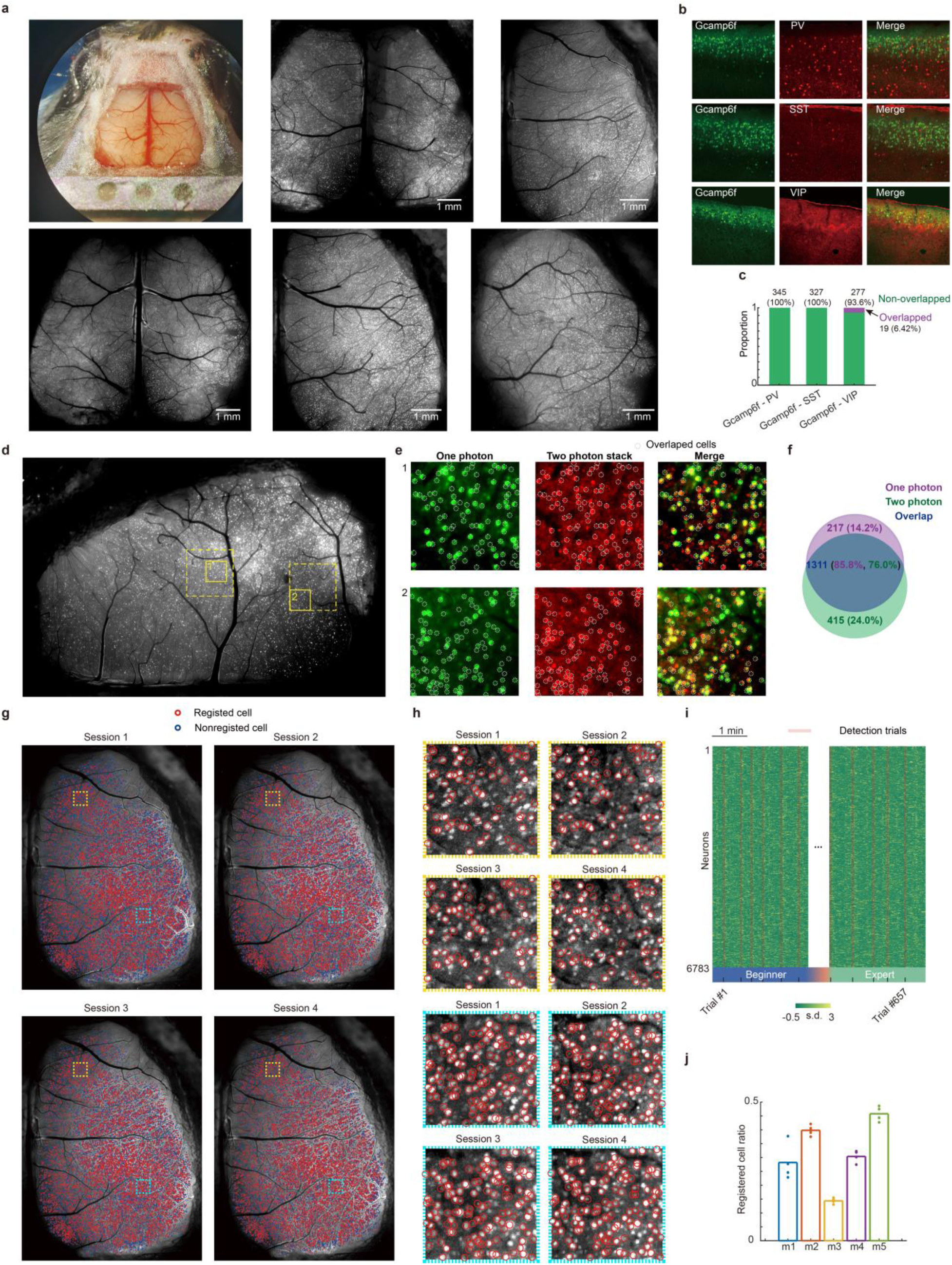
| Two photon verification of single neurons resolved using widefield imaging and neuronal registration across sessions. **a.** Surgical preparation and dataset overview. Top left, mouse with dorsal cortex craniotomy. Remaining panels, pixelwise standard deviation images from cortex-wide recordings across the 5 mice included in widefield experiments. The bottom middle panel is reproduced from Fig. 1f. **b.** Immunostaining to check co-localization between Gcamp6f and PV (top panels), Gcamp6f and SST (middle panels), Gcamp6f and VIP (bottom panels). **c.** Proportions of Gcamp6f expressing neurons in Rasgrf2 xAi148D co-expressing PV, SST or VIP. None of these neurons co-express PV or SST, only a small fraction (6.42%) co-express VIP. **d.** Fluorescence showing L2/3 neurons (pixel-wise standard deviation) revealed by widefield imaging. Yellow dashed rectangles mark regions of interest (ROIs) for two photon validation. **e.** Side-by-side comparison of widefield and two-photon imaging in the marked ROIs within solid boxes. Left panels, widefield imaging. Middle, two-photon imaging showing maximum intensity projections from 24-28 z-stacks (10 µm axial steps). Right, merged images. **f.** Widefield image identified 1,528 cells, comparable to the 1,726 cells identified with two photon imaging. About 86% of widefield identified neurons were resolved within the two-photon volumes. **g.** Example segmentations from sessions 1-4. **h.** Two zoom in view of ROIs marked in **g**, showing chronically tracked neurons. **i.** Heatmap of neural activity for registered cells (n = 6,783) from trial 1 to trial 657. **j.** The ratio of registered cells over extracted cells for five mice (m1-m5). Each dot represents a single session.

**Extended Data Fig. 3.**
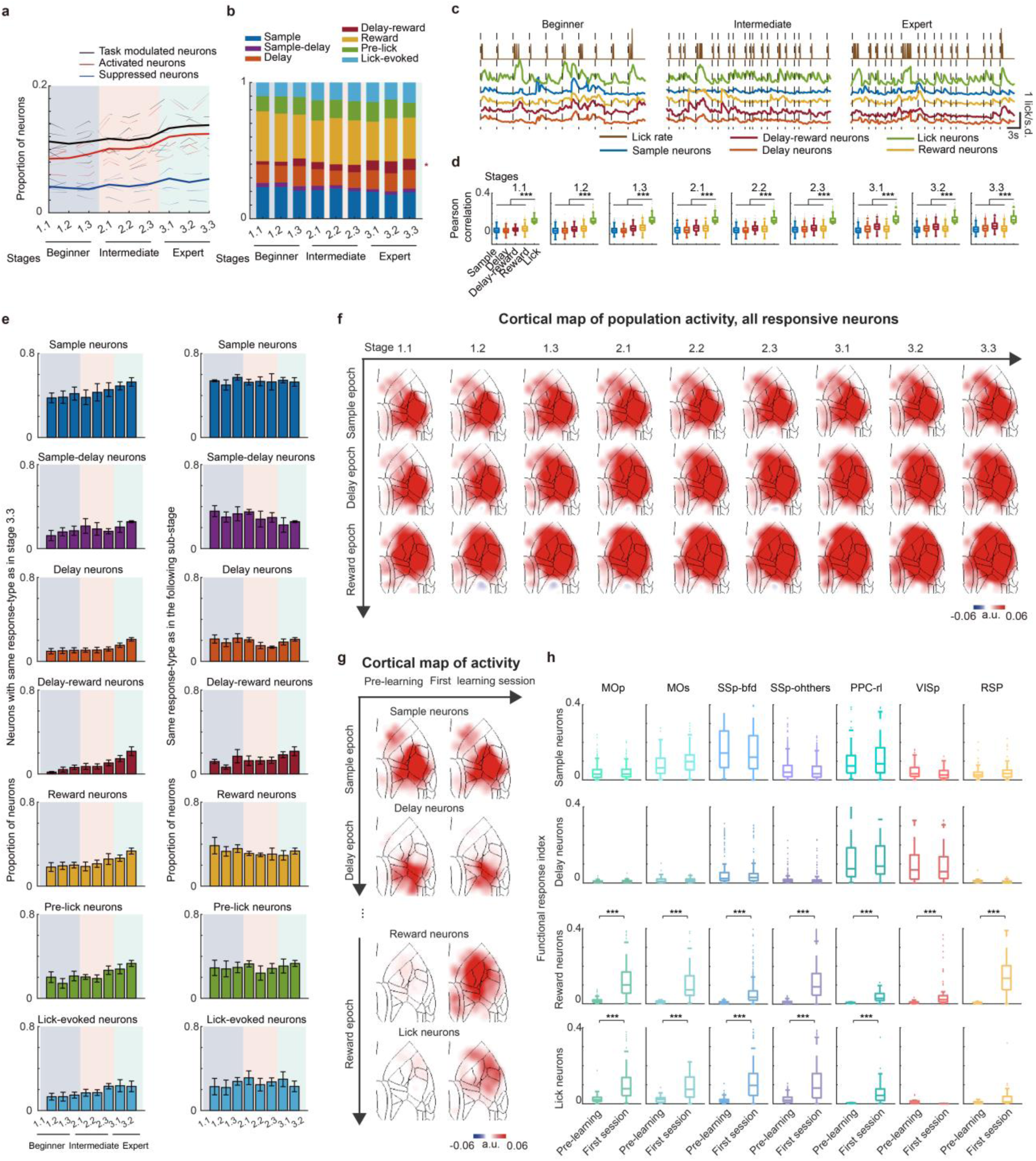
| Chronically imaged neurons across learning stages and cortical distribution of activity for responsive neurons. **a.** The proportion of modulated, activated and suppressed neurons across stages (chronically tracked neurons across mice, n = 28,904). The solid line represents the mean of 5 mice, and the dashed lines represent individual mice. **b.** Proportion of sample, sample-delay, delay, delay-reward, reward, pre-lick, lick-evoked neurons across learning. There is no sample-reward or sample-delay-reward neurons. Only delay-reward neurons increased significantly with learning. (\**P* < 0.05, Kruskal-Wallis ANOVA test). **c.** Lick and activity traces. Top, lick rate. Bottom traces, calcium activity during the inter trial interval for each response class. Dotted vertical lines indicate onset of licking bouts. Average activity of 5 classes of neurons from an example animal is shown. For the left panel, part of traces is reproduced from Fig. 2b. **d.** Quantification of the Pearson correlation coefficient for each type of neurons across learning stages. Lick neurons have significantly higher coefficients (\*\*\**P* < 0.001, Kruskal-Wallis ANOVA test, followed by a post hoc Dunn’s multiple comparison test). Box plot shows the median, upper and lower quartiles. Whiskers extending from the box indicating the upper and lower range. Points, outliers. **e.** Stability of response categories across stages. Left: proportion of neurons whose response class matches their stage 3.3 assignment. Similarity tends to increase for temporally adjacent stages. Right: proportion matching the subsequent stage. Error bar, SEM. **f.** The cortical map of neural activity across learning stages (columns) and trial epochs (rows). For each map, activity was averaged within the epoch over all responsive neurons. Overall spatial patterns remain largely stable across learning. **g.** Comparison between naïve (pre-learning) and beginner (the first learning session) mice. There is no evident change for the sample or delay activity. Reward and lick neurons increase their activity during the first learning session. **h.** Quantification of activity increase for sample, delay, reward and lick neurons (\*\*\**P* < 0.001, two-sided Wilcoxon rank sum test).

**Extended Data Fig. 4.**
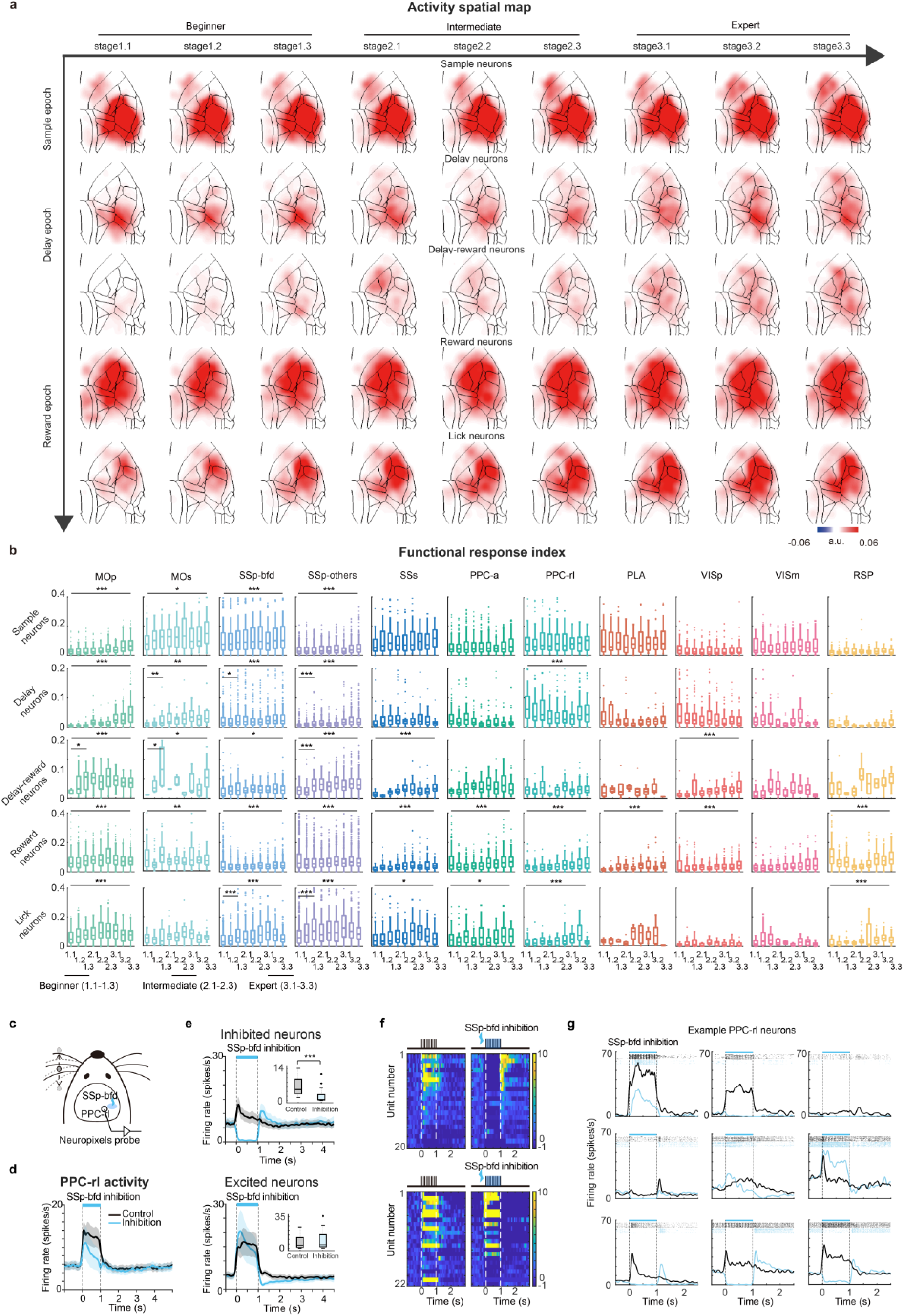
| The spatial distribution of response-type activity across learning stages and SSp-bfd effect on PPC-rl activity. **a**. Cortical maps of response-type activity across learning stages. Each row represents one response type. Values are mean activity within the indicated epoch, spatially smoothed with a kernel of 10 CCF pixels (262 µm). Data are from 5 mice. Same data as in Fig. 2 but with learning sub-stages. Delay, delay-reward, and lick neurons show enhanced activity across sensorimotor areas with leaning. PPC-rl shows reduced activity from stage 1 to stage 2. **b**. Quantification of the amplitude by response-type, learning stage and area (\**P* < 0.05, \*\**P* < 0.01, \*\*\**P* < 0.001, Kruskal-Wallis ANOVA test). Quantifications were performed to detect changes across the learning substages (from 1.1 to 3.3) or across the substages in beginner mice (substages 1.1, 1.2, and 1.3). Box plot shows the median, upper and lower quartiles. Whiskers extending from the box indicating the upper and lower range. Points, outliers. **c**. Schematic of recording from PPC-rl during whisker stimulation. Optogenetic inhibition of SSp-bfd was applied on 50% of interleaved trials. All whiskers were kept for this experiment. **d**. Average firing rate of PPC-rl neurons (n = 44). SSp-bfd inhibition reduces whisker evoked PPC-rl activity. Mean ± SEM. **e**. Subpopulation analysis by delay-epoch modulation. Top, neurons with delay activity (1 – 1.5 s) suppressed following SSp-bfd sample-epoch inhibition (compared with control, n = 20 neurons). Bottom, neurons with delay activity increased (n = 22). Mean ± SEM. Inset, quantification of PPC-rl neural activity within the first half of the delay epoch (1 – 1.5s). ***P < 0.001, two-sided test. **f**. Heatmaps of PPC-rl neural activity for the same set of neurons in **e**. Left, control trials without SSp-bfd inhibition; right, trials with inhibition. Top, neurons with suppressed delay activity. Bottom, neurons with increased delay activity. **g**. Example PPC-rl neurons illustrating the effects of SSp-bfd inhibition on whisker-evoked sample– and delay-epoch responses.

**Extended Data Fig. 5.**
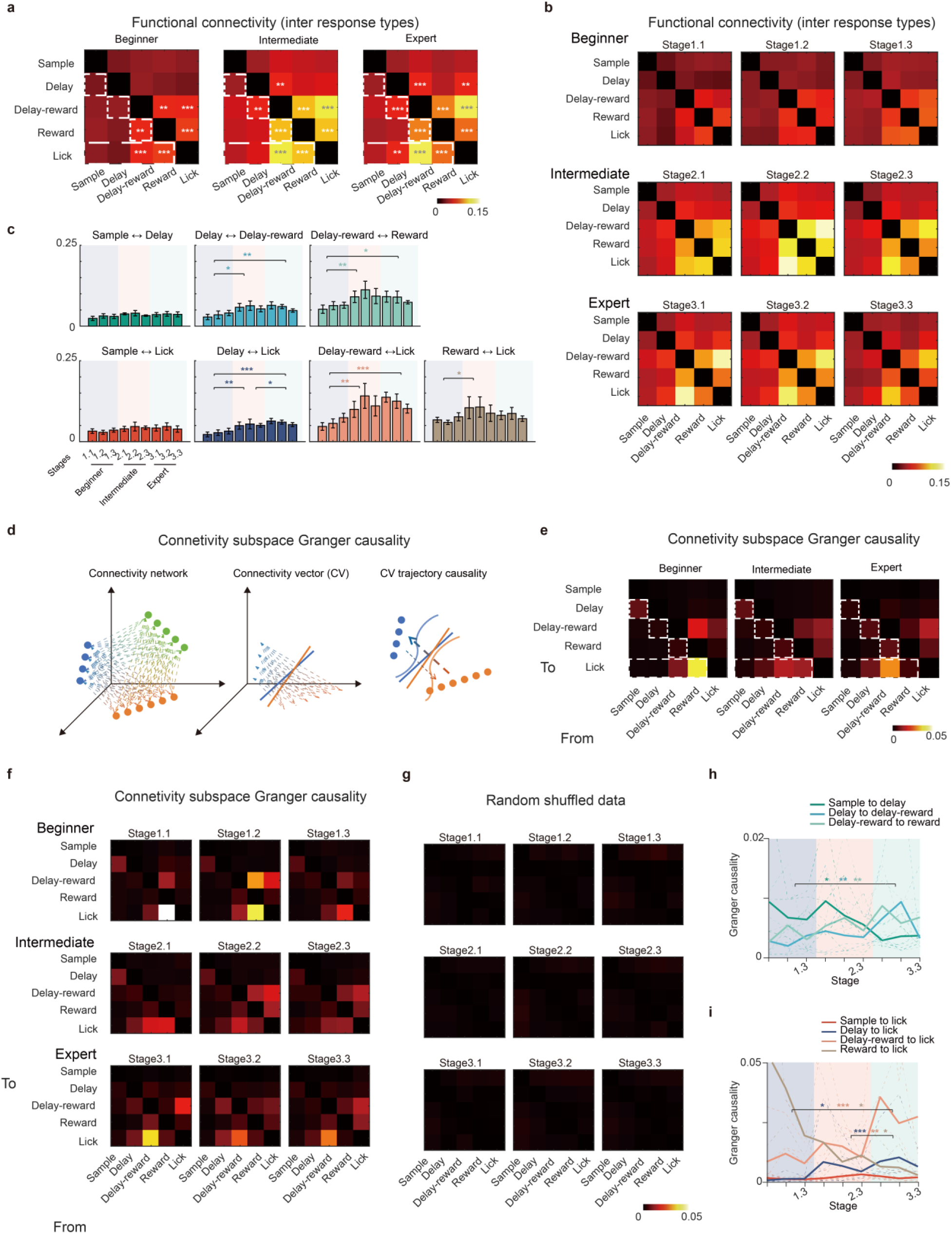
| Functional connectivity and Granger causality analyses among neuronal response groups across learning. **a.** Functional connectivity (FC) matrices for beginner, intermediate and expert mice. Heatmaps reflect noise correlations between response groups (835,441,216 neuron pairs from 5 mice). Learning strengthens FC of delay ↔ lick, delay-reward ↔ lick, reward ↔ lick, delay ↔ delay-reward, delay-reward ↔ reward. These pairs are significant higher than the global average across learning stages (\**P* < 0.05, \*\**P* < 0.01, \*\*\**P* < 0.001, two-sided Wilcoxon sign rank test). White rectangles highlight direct influence from different response groups to lick neurons, or along the temporal chains (see maintext). **b.** FC matrices for learning sub-stages. Same format as in **a**. **c.** Quantification of FC for temporal chain pairs (sample ↔ delay, delay ↔ delay-reward, delay-reward ↔ reward) and between each functional group and lick neurons (sample ↔ lick, delay ↔ lick, delay-reward ↔ lick, reward ↔ lick) across learning stages (*P < 0.05, **P < 0.01, ***P < 0.001, Kruskal-Wallis ANOVA test for stage averaged FC, followed by a post hoc Dunn’s multiple comparison test). Mean ± SEM. **d.** Schematic for Granger causality analysis. As we have thousands of simultaneously imaged neurons, Granger analysis of pairwise temporal sequence will be time consuming. We thus first reconstructed a connectivity matrix between one response type of neurons (for example, neurons with blue color) and the other response type of neurons (orange). The value of each element represents the connectivity estimated using resting-stage spontaneous activity during inter trial interval. We then performed principal components analysis (PCA) to identify the top components that explain 40% of the variance. After identifying the principal directions, we projected neural activity onto these components. We then performed pairwise Granger analysis of PCA projected time series and weighted component-wise results by their PCA loadings (see Methods). **e.** Granger-based functional connectivity (FC) matrices for beginner, intermediate and expert stages. Heatmaps reflect directed influence from source to target response groups, with higher values indicate stronger causal influence. Learning strengthens directed influence for delay → lick, delay-reward → lick, lick → reward, delay → delay-reward, delay-reward → reward. White rectangles highlight direct influence from different response groups to lick neurons, or along the temporal chains. **f.** Same as **e** but for learning sub-stages. **g.** Granger causality using randomly shuffled data did not reveal significant influence among response groups. **h.** Directed influence along the temporal chain across learning sub-stages. Directed influence increase for delay → delay-reward and delay-reward → reward while decreases for sample → delay (*P < 0.05, **P < 0.01, ***P < 0.001, Kruskal-Wallis ANOVA test, followed by a post hoc Dunn’s multiple comparison test). The solid line represents the mean of 5 mice, and the dashed lines represent individual mice. **i.** Directed influence onto lick neurons (sample → lick, delay → lick, delay-reward → lick and reward → lick). Directed influence increases for delay → lick and delay-reward → lick while decreases for reward → lick (*P < 0.05, **P < 0.01, ***P < 0.001, Kruskal-Wallis ANOVA test, followed by a post hoc Dunn’s multiple comparison test). The decrease is consistent with the observation that mice initially licked after reward delivery and then gradually switched to actively lick in preparation for reward.

**Extended Data Fig. 6.**
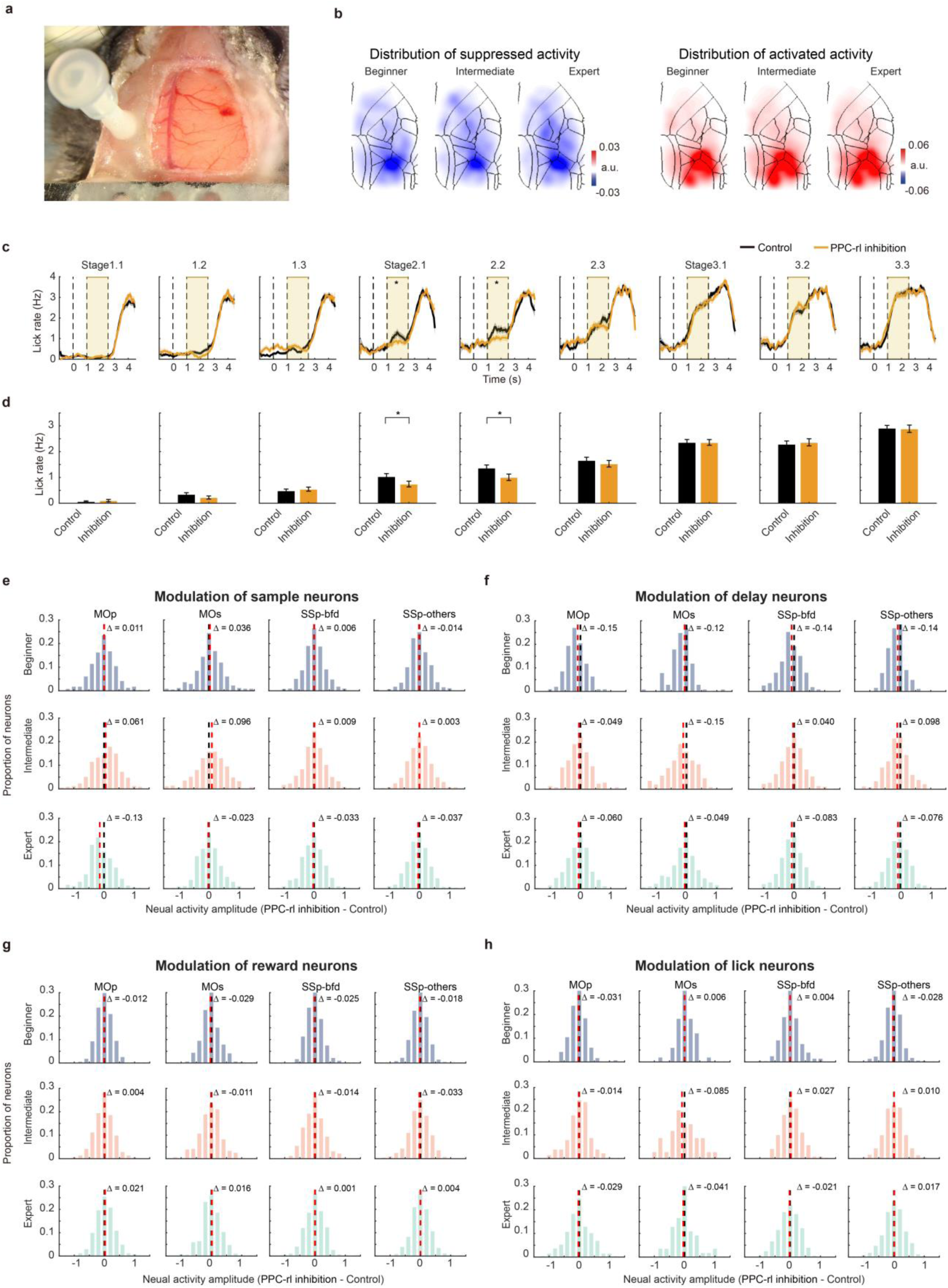
| Effects of PPC-rl inhibition on preparatory licking and on activity of different neuronal response types. **a.** Surgical preparation. Photograph of the craniotomy and optical fiber implantation. **b.** Spatial distribution of optogenetically modulated neurons. The left three panels, distribution of suppressed activity. The right three panels, distribution of increased activity. Neurons in PPC-rl are likely directly inhibited with light illumination and can show indirect activation via disinhibition. Neurons in sensorimotor areas are indirectly modulated. **c.** PPC-rl inhibition reduces preparatory licking during intermediate sub-stages 2.1 and 2.2 (\**P* < 0.05, two-sided Wilcoxon rank sum test, n = 3 mice, Related to **d**). Shading, Mean ± SEM. **d.** Quantification of inhibition effects on preparatory lick rate across learning stages **e.** Sample neurons. PPC-rl inhibition has minimal effect. Red dashed lines, mean activity during inhibition. Black dashed, zero. **f.** Delay neurons. PPC-rl inhibition produces mild suppression. **g.** Reward neurons showing minimal modulation. **h.** Lick neurons showing minimal modulation.

**Extended Data Fig. 7.**
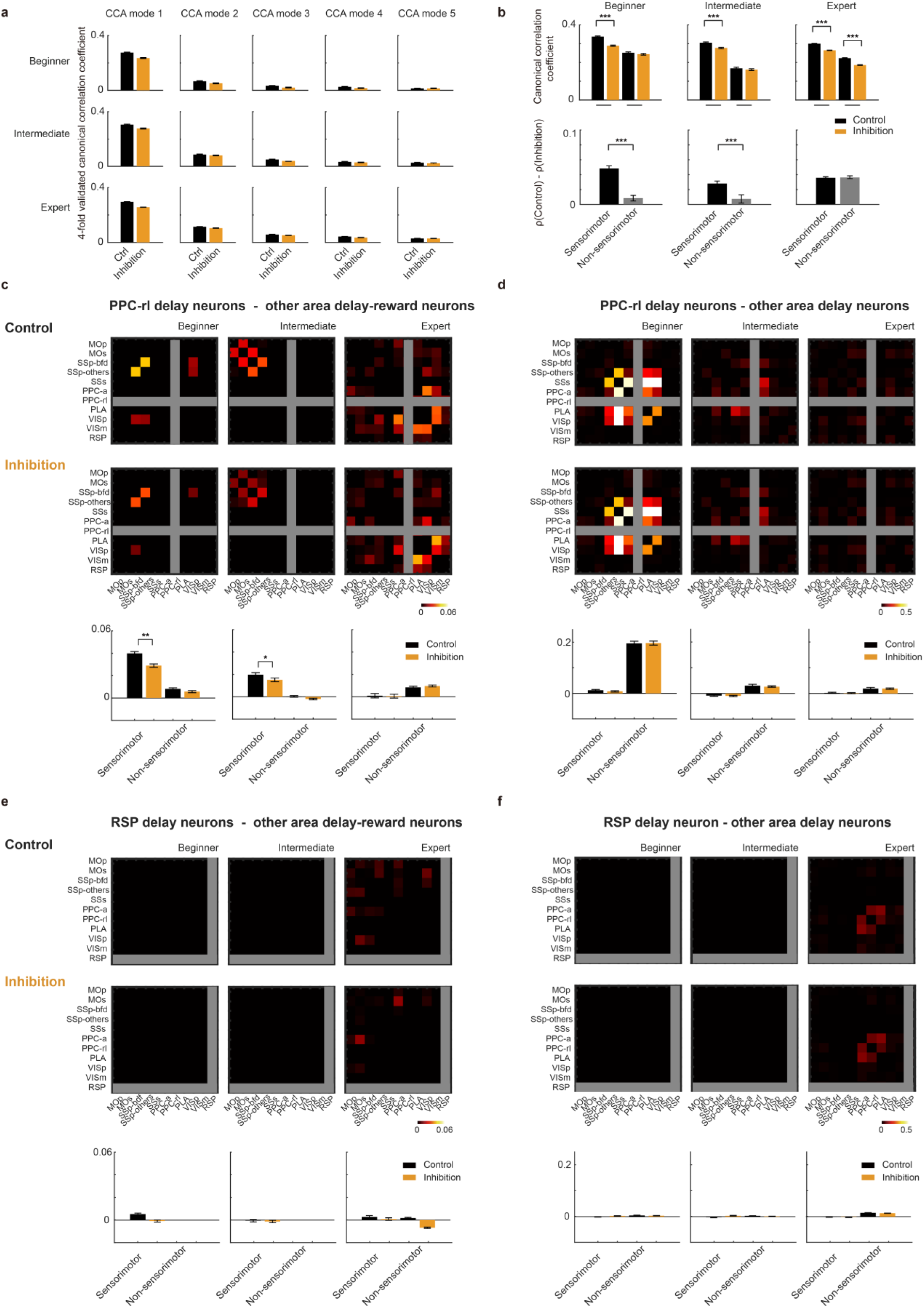
| Effects of PPC-rl inhibition on subspace communication. **a.** Canonical correlation of neural activity projected along each canonical correlation analysis (CCA) mode. CCA identifies co-fluctuation subspaces linking PPC-rl delay activity with delay-reward activity in each cortical area. For each CCA mode, we computed the correlation between PPC-rl delay activity and area-specific delay-reward activity and averaged across areas. The coefficients are relatively low as the delay activity and delay-reward activity are only partly overlap in time. Inhibition of PPC-rl mainly reduced the first CCA mode. Mean ± SEM. **b.** Inhibition effects by area class. Top: effects on the first CCA mode between PPC-rl and sensorimotor areas vs non-sensorimotor areas across learning stages (sensorimotor vs. non-sensorimotor, \*\*\**P* < 0.001, two-sided permutation test). Bottom: difference of inhibition effects between the two classes. Mean ± SEM. **C.** Heatmap matrix shows correlation coefficients for the first CCA mode between target-area pairs (indicated along x and y axes). CCA mode identifies low-dimensional communication subspace between source neurons (PPC-rl delay neurons) and individual target neurons (other area delay-reward neurons). Bright colors indicate stronger shared co-fluctuation with the source. Top, control across learning stages. Middle, PPC-rl inhibition. Bottom, quantification of CCA-derived coupling across area pairs and learning stages. PPC-rl inhibition reduced a common co-fluctuation mode within sensorimotor areas (\**P* < 0.05, \*\**P* < 0.01, two-sided permutation test). Mean ± SEM. **D.** Same format as **c** but for PPC-rl delay (source) to delay (target) neurons in each of the other areas. PPC-rl inhibition did not affect their co-fluctuations. **E.** Same format as **c** but for RSP delay (source) and delay-reward (target) neurons in each of the other areas. RSP inhibition did not affect their co-fluctuations. **F.** Same format as **c** but for RSP delay (source) and delay (target) neurons in each of the other areas. RSP inhibition did not affect their co-fluctuations.

**Extended Data Fig. 8.**
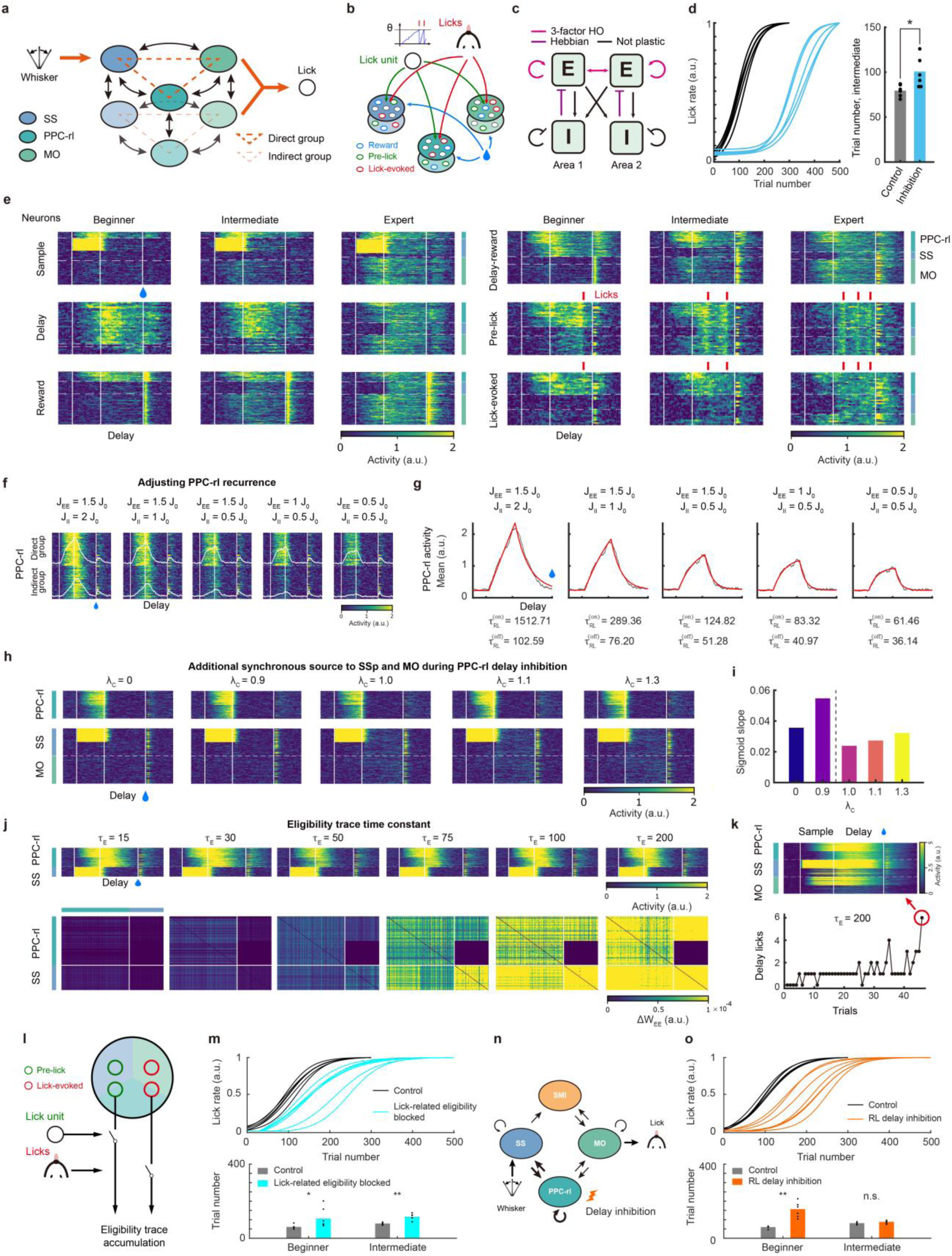
| Simulation of network dynamics under different parameterizations. **a.** Network architecture. The model comprises direct and indirect neuronal groups within SS, PPC-rl and MO. The SS direct group receives whisker input directly and forms long-range projections with the direct groups in PPC-rl and MO. Indirect groups (in PPC-rl and MO) do not connect to the SS direct group (by definition), but form other within-or across-area connections. **b.** Neurons modulated by reward, lick units (that drive licking) and lick movements are defined as “reward”, “pre-lick” and “lick-evoked” units, occupying 50%, 15%, 15% of neurons, respectively. **c.** Plasticity specification. E-to-E connections and local I-to-E connections are plastic (see Methods for details). 3-factor HO: three-factor Hebbian learning rule for Oja’s variant. **d.** Network simulation under control and PPC-rl delay inhibition conditions. We ran simulation 6 times as the results are stable. The intermediate learning stage with inhibition is also longer compared with control (\**P* < 0.05, two-sided Wilcoxon rank sum test), but the difference is much smaller compared with the beginner stage (24.95% vs 363.39%). **e.** Heatmaps of single trials. Representative activity from 6 functional groups across three areas illustrates sample, delay, delay-reward, reward, pre-lick and lick-evoked responses. For each heatmap, each row represents a single trial from one neuron. **f.** PPC-rl recurrence manipulations. Direct and indirect group activities on the first trial for different recurrence settings. White traces are overlaid on each panel to show population-averaged activity. J_EE_ and J_II_, average PPC-rl E-to-E and I-to-I strengths. J_0_, default average connection strength, equal to mean of a mean-shifted standard normal distribution (μ = 0.2), clipped at 0.3. **g.** Quantifying PPC-rl network recurrence by calculating response time constants of averaged PPC-rl activity in the first trial (see Method for details). Average involves both direct and indirect group (gray line). Red line shows the fitted response. Onset time constant (*τ*^(*on*)^) are typically larger than offset time constant (*τ*^(*off*)^) as the former one involves synaptic communication delay. These two values are positively correlated, and using *τ*^(*off*)^ yields the same conclusion as in Fig. 6h. **h.** PPC-rl delay inhibition and the rescue using a low-dimensional driving source to sensorimotor areas. Injecting an additional co-fluctuation into SS and MO during the delay (strength *λ_C_* ≥ 1) restores delay activity and accelerates weight changes as *λ_C_* increases. **i.** Learning trajectory is qualitatively different under *λ_C_* ≤ 0.9 and *λ_C_* > 1, implying that effective learning during PPC-rl inhibition requires surpassing a critical threshold of sensorimotor co-fluctuation. **j.** Effect of eligibility trace duration (unit, time step). First-trial activities of PPC-rl and SS direct-group neurons (top) and corresponding synaptic updates (bottom) for different eligibility-trace time constants (*τ_e_*). **k.** Unstable learning and epileptic activity when *τ_e_* is excessively large (*τ_e_* = 200 time steps). **l.** Lick unit elicits preparatory related activities for pre-lick neurons while licking events modulate lick-evoked neurons. We blocked the eligibility traces generated by these two types of lick-related activities to test the effect of preparatory licking for learning. **m.** Learning trajectory and learning stage comparison between control condition and the eligibility-blocked condition (\**P* < 0.05, ** *P* < 0.01, two-sided Wilcoxon rank sum test). **n.** Schematic of the four-area model, where SS does not directly communicate with MO. Instead, a sensorimotor interface (SMI) region is introduced to relay SS activities to MO, and also forming recurrent excitatory connections with MO, mimicking the cortico-striato-thalamocortical pathway. The function of PPC-rl is tested by inhibiting PPC-rl neurons during the delay epoch. **o.** Learning trajectory (top) and learning stage (bottom) comparison between the control and PPC-rl inhibition condition (** *P* < 0.01, two-sided Wilcoxon rank sum test). Inhibition of PPC-rl in the four-area model produced similar results as simulation using the three-area model.

**Extended Data Fig. 9.**
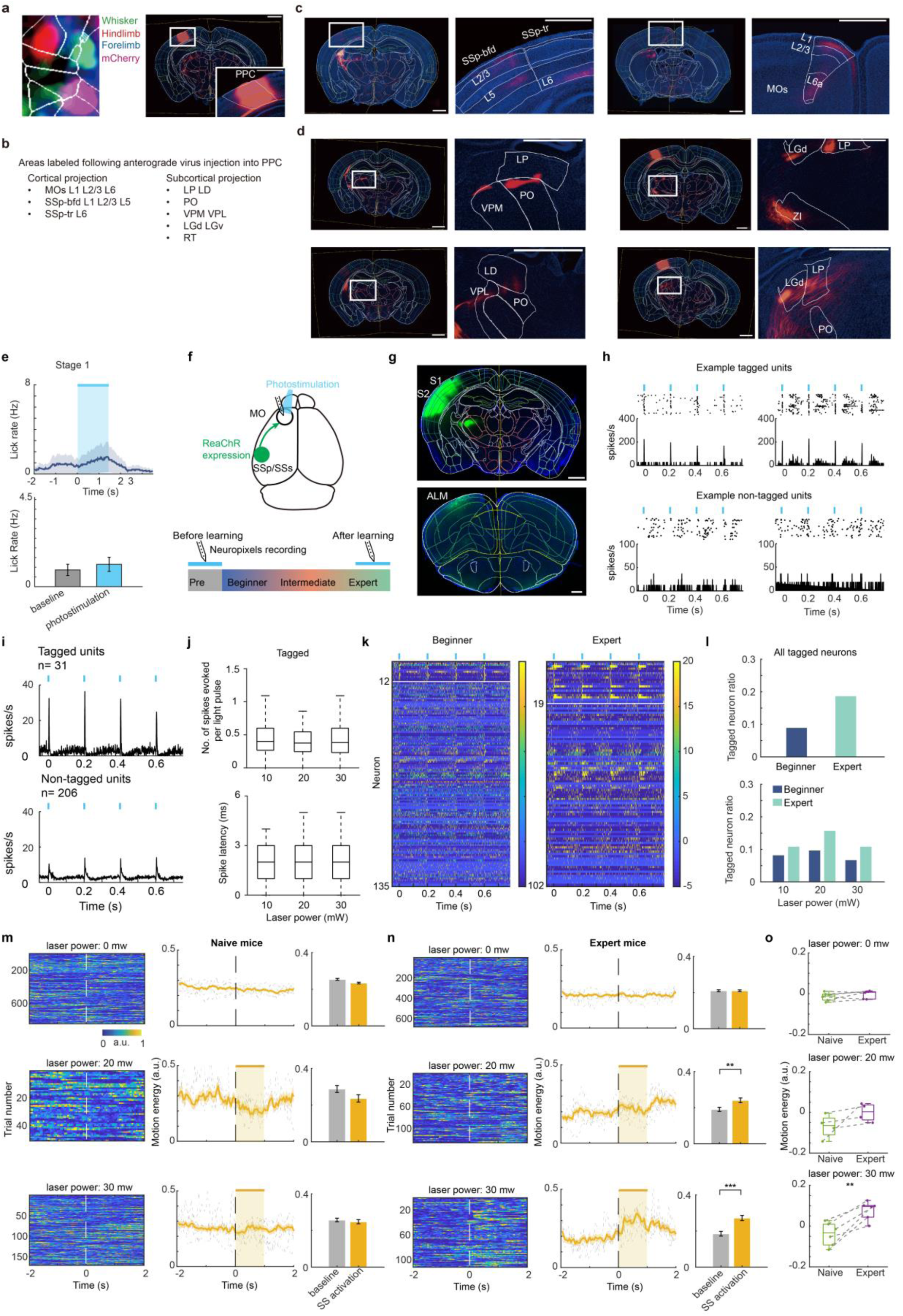
| PPC-rl projections in sensorimotor cortices, experimental validation of PPC-rl activation, and plasticity along the somatosensory-motor pathway, and. **a.** Viral targeting of PPC-rl via ISI with AAV2/9-hSyn-mCherry. Scale bar is 1 mm. **b.** List of areas that were labeled by the PPC-rl projection axons (based on injections in 3 mice). **c.** Example brain slices showing PPC-rl downstream cortical areas including SSp and MOs. Scale bar is 1 mm. **d.** Example brain slices showing PPC-rl downstream subcortical areas. Scale bar is 1 mm. **e.** Optogenetic activation of PPC-rl alone did not increase preparatory licking in beginner mice. However, when paired with whisker stimulation during training, PPC-rl activation accelerated learning. **f.** Schematic of long-range connectivity measurement from somatosensory cortex (SSp-bfd and SSs) to motor cortex (the anterior lateral motor cortex ALM which is the secondary motor area for licking) using optical tagging (see Methods). Optical tagging was performed in both naïve and expert mice. **g.** Top, ReaChR expression achieved by injecting rAAV-CaMKllá-Cre in the left SSp-bfd and SSs of CAG-LSL-ReaChR-mCit mice. Bottom, labeled axonal projections in ALM. **h.** Top, Example tagged ALM units exhibiting short-latency and reliable responses to photostimulation of somatosensory axons. Bottom, Example non-tagged ALM units showing suppressed or unresponsive activity profiles. **i.** Average light-evoked responses of tagged (n = 31) and non-tagged (n = 206) units. **j.** Response amplitude and spike latency of tagged units across laser powers (10, 20 and 30 mW). Box plot shows the median, upper and lower quartiles. Whiskers indicate data range. **k.** Heatmaps of all recorded ALM units aligned to photostimulation. White horizontal lines separate tagged and non-tagged populations. **l.** Proportion of optically tagged units in naïve and expert mice (top) and as a function of laser powers (bottom). **m.** Motion energy during photostimulation of somatosensory axons for naïve mice. Left, trial-by-trial heatmaps. Middle, mean ± SEM, and the dashed lines represent individual mice. Right, comparison of motion energy before and during photostimulation. Rows correspond to different laser powers. **n.** Same as I but for expert mice. Photostimulation significantly increased orofacial movements (** *P* < 0.01, *** *P* < 0.001, two-sided Wilcoxon rank sum test). **o.** Quantification of motion energy changes in naïve versus expert mice (** *P* < 0.01, two-sided Wilcoxon rank sum test).

